# Kidins220 regulates the development of B cells bearing the λ light chain

**DOI:** 10.1101/2022.04.04.486937

**Authors:** Anna-Maria Schaffer, Gina J. Fiala, Miriam Hils, Eriberto Noel Natali, Lmar Babrak, Laurenz A. Herr, Marta Rizzi, Enkelejda Miho, Wolfgang Schamel, Susana Minguet

**Author notes:** Corresponding author: Susana Minguet.

## Abstract

The ratio between Igκ and Igλ light chain (LC)-expressing B cells varies considerably between species. We recently identified Kinase D-interacting substrate of 220 kDa (Kidins220) as an interaction partner of the BCR. *In vivo* ablation of Kidins220 in B cells resulted in a marked reduction of λLC-expressing B cells. Kidins220 knockout B cells fail to open and recombine the genes of the *λLC* locus, even in genetic scenarios where the *κLC* genes cannot be rearranged or where the κLC confers autoreactivity. *κLC* gene recombination and expression in Kidins220-deficient B cells is normal. Kidins220 regulates the development of λLC B cells by enhancing the survival of developing B cells and thereby extending the time-window in which the *λLC* locus opens and the genes are rearranged and transcribed. Further, our data suggest that Kidins220 guarantees optimal pre-BCR and BCR signaling to induce *λLC* locus opening and gene recombination during B cell development and receptor editing.

**One Sentence Summary:** We demonstrate that the scaffold protein Kidins220 regulates the development of λLC B cells by supporting B cell precursor survival and optimizing pre-BCR and BCR signaling to open, recombine, and transcribe the genes of the *λLC* locus.

## INTRODUCTION

Antigen recognition in B cells is mediated by the B cell antigen receptor (BCR), composed of two immunoglobulin (Ig) heavy chains (HC) and two Igκ or Igλ light chains (LC). They form the BCR complex together with the associated Igα and Igβ heterodimer to transmit signals for B cell development, proliferation, survival, and activation. Each B cell expresses a BCR with a given specificity, which is acquired by a progressive rearrangement of the *variable* (V), *joining* (J) and, in case of the HC, *diversity* (D) gene segments of the BCR *HC* and *LC* loci in the bone marrow (BM) (reviewed in (*1*, *2*)).The key enzymes that facilitate V(D)J recombination are recombination-activating gene 1 (RAG1) and RAG2 proteins (*3*, *4*). V(D)J recombination is initiated at the pro-B cell stage (*5*). An in-frame rearrangement at the *IgHC* genes leads to the expression of a μHC that is binding to the λ5 and VpreB components of the surrogate light chain (*6*–*8*). Together with the Igα/β signaling subunits these chains form the pre-BCR complex, which is first expressed on large pre-B cells. Signals from the pre-BCR and the interleukin (IL)-7 receptor induce a proliferative burst that is followed by cell cycle attenuation, promoting *IgLC* V- to J-gene segment rearrangement in the small pre-B cell stage (*8*–*13*). A productive rearrangement leads to the pairing of the pre-existing HC with the newly generated LC, forming the IgM-BCR expressed first on the immature B cells in the BM (reviewed in (*2*, *14*)).

*LC* locus opening and recombination during B cell development and receptor editing (a process changing the specificity of the BCR by secondary *IgLC* VJ-gene rearrangements) is dependent on transcription factors including Ikaros, Aiolos, interferon regulatory factor (IRF)-4, IRF-8 and E2A (*11*, *15*–*25*). Genetic mouse models individually lacking some of these transcription factors show impaired *LC* locus opening, affecting λLC expression more severely (*18*, *21*, *22*, *25*). The activity of these transcription factors is directly or indirectly regulated by pre-BCR and BCR signaling. Mice lacking a signaling competent pre-BCR or BCR, or lacking signaling molecules downstream of the pre-BCR or BCR including the adapter protein SLP65, Bruton’s tyrosine kinase (BTK) and phospholipase Cγ2 (PLCγ2), are characterized by an altered LC expression (*15*, *26*–*29*). Specifically, mice deficient in SLP65 show reduced *LC* germline transcripts especially from the λLC (*16*, *26*, *28*). Likewise, the absence of BTK, a kinase that is recruited by SLP65, severely reduces *λLC* germline transcripts and LC expression (*16*, *26*, *27*). The concurrent ablation of both SLP65 and BTK abrogates *κLC* and *λLC* germline transcription and BCR surface expression (*16*, *26*). PLCγ2 is recruited to SLP65, regulating the calcium signaling downstream of the pre-BCR and BCR (*29*, *30*). PLCγ2-knockout (KO) mice show a strong reduction of λLC^+^ B cells and a mild reduction of *κLC* germline transcripts (*15*, *29*). The combined deficiency of BTK and PLCγ2 almost completely abrogated LC expression (*29*).

Pre-BCR signaling is important for the induction of *IgLC* VJ-gene rearrangement at both *LC* loci (κ and λ). However, the ratio between the usage of the two LCs diverges greatly among species (*31*) and the regulation of their differential expression is still not fully understood. The primary B cell repertoire in mice is dominated by the κLC that is roughly ten to 20 times more frequent than the λLC (*31*, *32*). It is generally accepted that λLC B cell generation is favored when rearrangement at the *κLC* locus is unsuccessful, or when the immature κLC-containing BCR confers autoreactivity (*33*, *34*). In the latter case, the BCR specificity is modified by receptor editing. The ratio of κ/λ LC is further impacted by the survival of developing B cells. Extending the life-span of B cell precursors by overexpression of the anti-apoptotic protein B cell lymphoma 2 (BCL2), promotes the generation of λLC B cells (*27*, *35*, *36*). In line with this, limiting B cell survival by genetically abrogating the NF-κB signaling pathway, results in mice with a reduced amount of λLC B cells (*36*). Based on these observations, the generation of λLC B cells mainly depends on (i) the ability and kinetics of the *λLC* locus opening, (ii) the life-span of developing B cells, and (iii) the efficiency of receptor editing during tolerance induction (*36*).

In this study, we investigated the differential regulation of the κ-*versus* λLC expression. We have identified the transmembrane protein Kidins220 as a new binding partner of the BCR (*37*). Kidins220 was first described as Kinase D-interacting substrate of 220 kDa in neuronal cells (*38*, *39*). As a scaffold protein, Kidins220 is implicated in multiple cellular processes like survival, proliferation and receptor signaling, among which are also the antigen receptors of T- and B cells (*37*, *40*, *41*). The complete genetic deletion of Kidins220 is embryonically lethal (*42*). Conditional mb1Cre-mediated B cell specific Kidins220 KO mice (B-KO) showed reduced BCR signaling, and almost complete loss of λLC B cells in the BM and periphery, with only mild effects on the κLC compartment (*37*). Our new findings presented here indicate that Kidins220 is crucial for the generation of λLC B cells by supporting B cell progenitor survival and pre-BCR and BCR signaling, which allows for *λLC* locus opening, recombination and protein expression.

## RESULTS

### Subhead 1: Kidins220 B-KO mice show a skewed primary BCR repertoire

Despite almost normal B cell numbers, B cell specific Kidins220 KO (B-KO) mice show an approximately 75% reduction of B cells carrying a λLC-containing IgM-BCR on the cell surface (**Fig. 1A**) (*37*). To further understand the molecular mechanism leading to this phenotype, we performed in-depth analysis of the primary IgM-BCR repertoire of control (CTRL) and B-KO mice. We FACS-sorted immature (B220^+^IgM^+^IgD^-^) B cells from the BM of a pool of three individual mice per genotype and performed sequencing analyzing paired (HC and LC), full-length V(D)J sequences from cDNA of single cells. Successful purification of immature B cells was confirmed by the annotation of 98-99% of HCs to the IgM isotype (**Fig. 1B**, left). In the CTRL, 83% of all μHCs were co-expressed with the κLC and the remaining 17% with various subclasses of the λLC (**Fig. 1B**, right). In contrast, in B-KO B cells, the μHC was almost exclusively co-expressed with the κLC (98%) and only 2% of all cells contained a λLC. Thus, the absence of λLC^+^ BCRs on the surface of B cells in B-KO mice reflects absent production of mRNA of λLCs from all λLC subclasses.

**Fig. 1.**
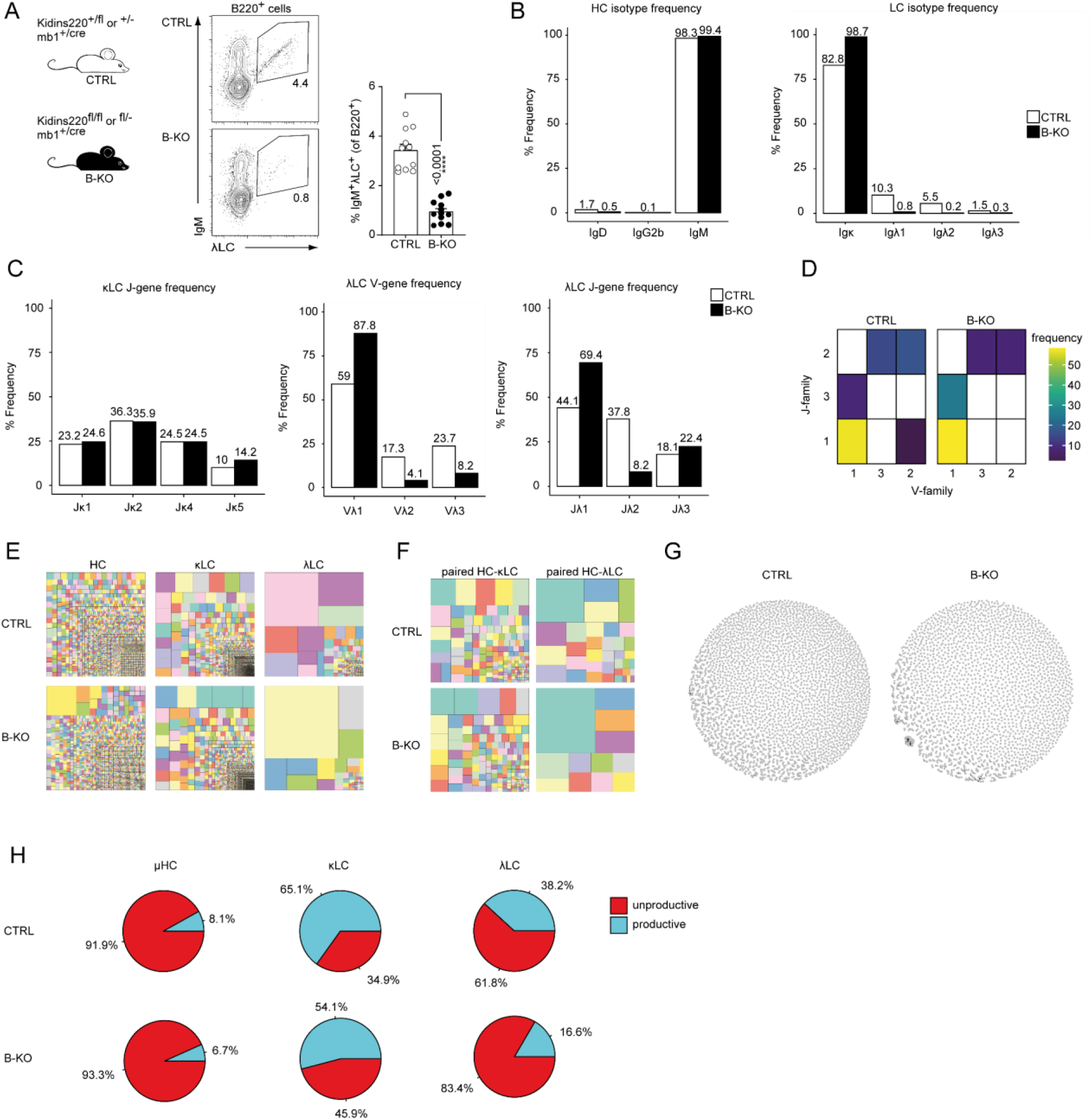
Kidins220 immature B-KO cells show a skewed primary BCR repertoire. (**A**) Schematics of the genotypes of the *Kidins220* locus and cre-recombinase expression for control (CTRL) and B cell specific knockout (B-KO) mice (left). Representative flow cytometric BM analysis using antibodies for B220, IgM and λLC as well as statistical quantification (right, five independent experiments were pooled; n = 11-12 mice per genotype) The mean ± SEM is plotted. Statistical analysis compared CTRL and B-KO samples using Mann-Whitney test. *P* value is indicated. (**B-H**) Immature B cells (B220^+^IgD^-^IgM^+^) of three individual mice per genotype were pooled and subjected to single cell sequencing analyzing full-length *Ig*-gene V(D)J recombination status and BCR repertoire based on cDNA. More than 10.000 cells per genotype were analyzed. Only productive recombinations are used for the analyses in (**B-G**). (**B**) The Frequencies of the individual HC and LCs isotypes in CTRL and B-KO of all obtained HC or LC sequences are plotted. (**C**) The Frequencies of *V-* and *J-* gene sequences in κLC and λLC of all obtained *V-* or *J-* gene sequences within the respective LC isotype are plotted. (**D**) Heatmap of the *V-J* gene combinations in the λLC. (**E**) Tree maps illustrating clonal CDR3 frequency of HC, κLC and λLC. Each square represents an individual CDR3, and its frequency correlates with the square size. (**F**) Tree maps illustrating the clonal CDR3 frequency of the whole BCR (paired HC-LC). Each square represents an individual CDR3, and its frequency correlates with the square size. (**G**) Primary antibody repertoire showing unique CDR3 sequences (nodes) and their similarity (edges) by Levenshtein distance (LD). LD = 1 represents one amino acid difference in the CDR3 sequence. (**H**) Productive and unproductive read frequencies.

We next investigated whether Kidins220 plays a role in the usage of specific *V-* and *J*-genes. The murine *IgHC* locus consists of more than 100 V*-*, 8-12 D- and 4 J-genes (*43*). We compared the use of each gene segment between CTRL and B-KO cells by performing a Pearson correlation test. We obtained very strong correlations for *IgHC* V- and J-gene usage (r = 0.97 and 0.98 respectively), suggesting that Kidins220 does not influence which V(D)J-gene is used for the μHC of the BCR (**Fig. S1A**). The murine *Igκ* locus consists of at least 101 functional V-genes as well as four functional J-genes (*44*) (**Fig. S1B**). Likewise, we observed a good correlation of the *Igκ* V- and J-gene usage (r = 0.96 and r = 1) between CTRL and Kidins220-deficient cells (**Fig. S1A**), confirming a rather intact κLC repertoire. Still, B-KO B cells showed a slightly increased use of *J_κ_5* (14.2%) when compared to CTRL (10%) (**Fig. 1C**). Use of *J_κ_5* correlates with increased secondary *Igκ* V- to J-gene rearrangements. It frequently occurs before deletion of part of the *κLC* locus via a recombining sequence recombination and opening of the *λLC* locus (*45*, *46*). The murine *Igλ* locus comprises one cassette including *V_λ_2* and *V_λ_3* upstream of *J_λ_2C_λ_2* (λ2), and a second cassette containing *V_λ_1* upstream of *J_λ_3C_λ_3* (λ3) and *J_λ_1C_λ_1* (λ1) (**Fig. S1B**) (*47*, *48*). In mature wild type B cells, the different λLCs are used at a frequency of 62%, 31% and 7% for λ1, λ2 and λ3, respectively (*48*, *49*). This pattern was confirmed by our sequencing results for immature B cells in CTRL mice (**Fig. 1B,**right). The *Igλ* V-gene usage showed a preference for *V_λ_1* over *V_λ_3* over *V_λ_2*, independent of the presence of Kidins220 (**Fig. 1C**). However, Kidins220-deficient B cells showed a relative increase of *V_λ_1*-gene usage (88%*versus* 59% in CTRL) (**Fig. 1C**). Both CTRL and B-KO immature B cells preferentially used the *J_λ_1*-gene segment (44% and 69%, respectively). However, B-KO B cells used *J_λ_3* more frequently than *J_λ_2*. These observations suggest that in the absence of Kidins220, the genes of the second λLC cassette are favored (**Fig. 1C**). Indeed, the Pearson correlation coefficient for the *Igλ* J-gene segments was weak (r = 0.51) (**Fig. S1A**). In the *λLC* locus, VJ-gene recombination preferentially takes place within the same cassette and is almost absent between cassettes (*50*, *51*). In line with this, we almost exclusively detected recombination of *V_λ_1* to either *J_λ_3* or *J_λ_1,* whereas *V_λ_2* and *V_λ_3* recombined to *J_λ_2* (**Fig. 1D**). Both CTRL and Kidins220-deficient λLC B cells showed a prominence of *V_λ_1-J_λ_1* joins. We next compared the HC, κLC and λLC repertoires of CTRL and B-KO immature B cells by analyzing the frequency of unique CDR3s in all chains. Both genotypes showed a clear polyclonal distribution for the κLC, but polyclonality was slightly reduced for the HC and strongly restricted for the λLC in the B-KO immature B cells (**Fig. 1E**). The analysis of paired HC-LC combinations highlighted that indeed the repertoire of those BCR bearing a λLC was strongly restricted in the B-KO immature B cells (**Fig. 1F**). We did not observe any predispositions for potentially autoreactive BCRs as characterized by longer or more positively charged CDR3s within their variable domain (**Fig. S1C**, **D**).

We further analyzed the architecture of the BCR repertoire by similarity networks connecting CDR3 nodes by edges when the CDR3 sequence differs by one amino acid (Levenshtein distance = 1) (*52*). Unexpectedly, the primary B cell repertoire of B-KO mice showed a specific clonal expansion, especially of one single clone containing the *V_H_10-1-gene* segment (**Fig. 1G**). The usage of this *IgHC* V-gene segment has been previously associated with anti-DNA antibodies and herpesvirus infections (*53*, *54*). Lastly, we determined the ratio between productive and unproductive rearrangements (**Fig. 1H**). Unproductive sequences are defined as out-of-frame sequences, sequences containing premature stop codons, orphon genes or non-IgHC sequences (*55*). They encompass a significant high proportion of the raw outputs (*55*) and are usually removed during a preprocessing step prior to data analysis (as done for **Fig. 1B-G**; **Fig. S1**). Both, CTRL and B-KO, showed similar frequencies of sequences defined as unproductive rearrangements (92% and 93%, respectively) from the *IgHC* locus. In contrast, the frequency of sequences defined as unproductive rearrangements from the *IgLC* loci was increased in B-KO compared to CTRL cells: 1.2 times for κLC, and 2 times for the λLC (**Fig. 1H**). Deeper investigations revealed that these unproductive rearrangements within the *λLC* locus were caused by a series of premature stop-codons (data not shown). Taken together, genetic ablation of Kidins220 in B cells skews the primary BCR repertoire mainly due to unsuccessful production of λLCs.

### Subhead 2: Kidins220 is essential for the opening of the *λLC* locus

Kidins220-deficient B cells failed to express λLC. The expression of the κLC and λLC depends on at least three factors: i) the ability and kinetics of the *κLC* and *λLC* loci opening, ii) the life-span of the pre-B cells and iii) the level of receptor editing during tolerance induction (*27*). Hence, we generated BM-derived pro-/pre-B cell cultures from CTRL and B-KO mice BM to study their ability to open the *LC* loci (**Fig. 2A**). After a seven-day culture in the presence of IL-7, we obtained a population almost homogeneous for surface expression of B220 and lacking a surface BCR, indicative of pro- and pre-B cells (data not shown (*56*)). Subsequent IL-7 withdrawal led to a relative increase of cells expressing a BCR on their surface (*27*, *57*). CTRL and B-KO B cells upregulated κLC^+^ IgM-BCR (**Fig. 2B**). The proportion of λLC^+^ IgM-BCRs was higher in CTRL compared to B-KO B cells at day 3 of culture, suggesting that the *in vivo* phenotype is B cell intrinsic.

**Fig. 2.**
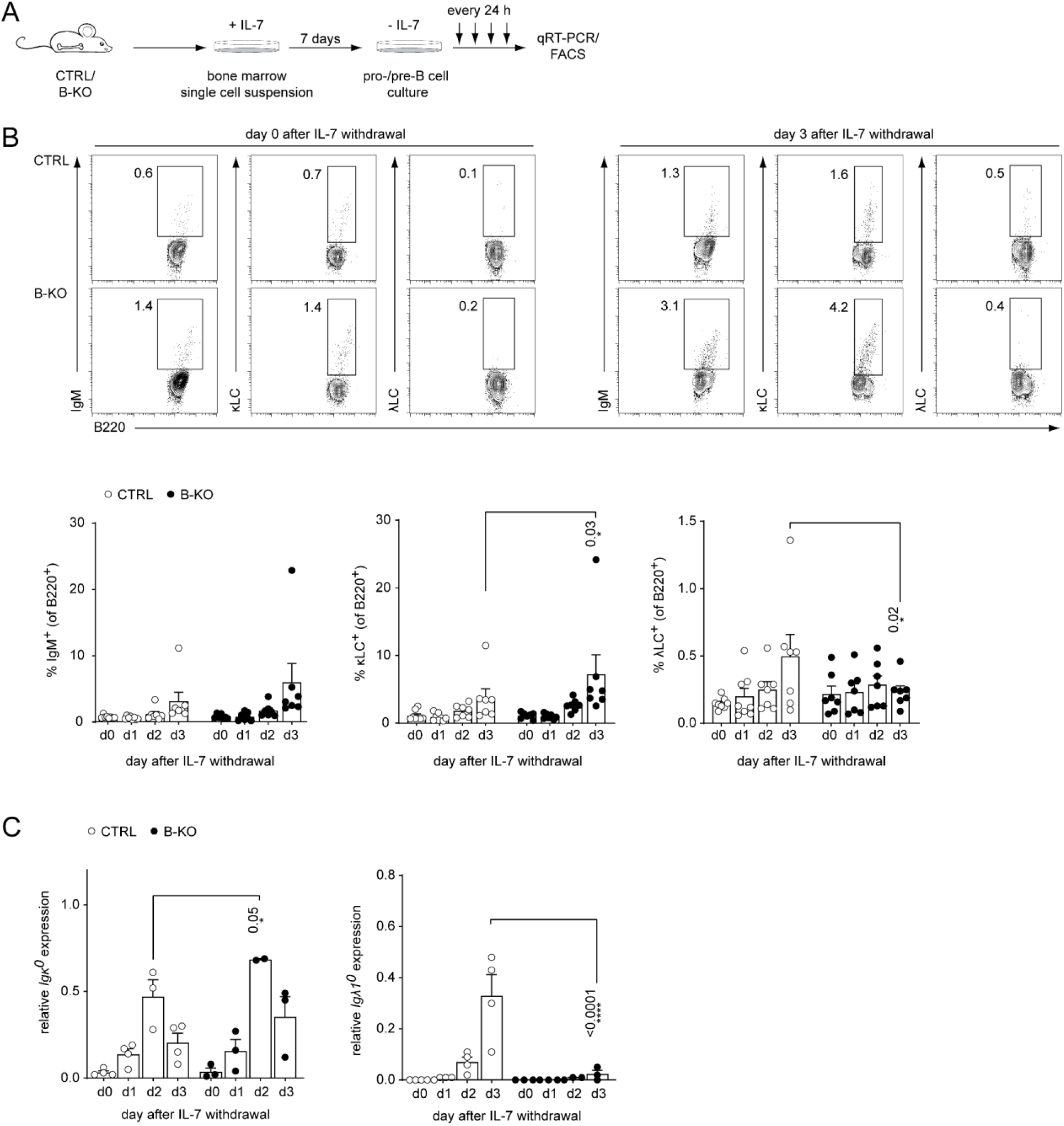
Kidins220 is required for the opening of the *λLC* locus in pro-/pre-B cells. (**A**) Experimental setup to generate primary BM derived pro-/pre-B cell cultures. Total BM was isolated from CTRL and B-KO mice and cultured for seven days in medium supplemented with IL-7. Next, IL-7 was removed, and samples were taken immediately (day 0) and every 24 hours for subsequent analyses. (**B**) Flow cytometry assayed surface expression of B220, IgM, λLC and κLC. Representative plots of day 0 and day 3 as well as statistical analyses are shown (n = 7-8 per genotype; each symbol represents one mouse). (**C**) Isolated RNA was reverse transcribed to assay the relative expression of germline transcripts of the *κLC* and *λLC* loci by qRT-PCR. Normalization was done using *Hrpt* (n = 2-4 per genotype; each symbol represents one mouse). In all graphs, the mean ± SEM is plotted. Statistical analysis compared CTRL and B-KO samples for each day using ANOVA test. Only *P* values < 0.05 are indicated.

Prior to *IgLC* VJ-gene rearrangement, the gene locus is opened and germline transcripts are produced (also known as sterile transcripts that do not encode for any protein but are proposed to possess regulatory functions) (*58*, *59*). We analyzed the relative expression of germline transcripts from the *κLC* and *λLC* loci in CTRL and B-KO pro-/pre-B cell cultures by qRT-PCR (**Fig. 2C, Fig. S2A**). The relative amount of *κLC* germline transcript and the kinetics of induction was similar between CTRL and B-KO B cells (**Fig. 2C**). However, Kidins220-deficient B cells failed to induce germline transcription from the *λLC* locus to the levels of CTRL cells (**Fig. 2C**). The *κLC* germline transcripts in the CTRL cells peaked at day two compared to day three for the *λLC* germline transcripts, in line with the findings that activation of the *κLC* locus precedes the *λLC* locus (*33*, *34*, *60*). LC germline transcription is dependent on the expression of E2A proteins (E12 and E47), which in turn regulate the RAG1 and RAG2 proteins responsible for VJ-gene recombination (*21*, *25*, *61*). We did not detect major differences in the expression of *E12, E47, Rag1* and *Rag2* mRNA as assayed by qRT-PCR between CTRL and B-KO B cells (**Fig. S2B).** Together, these data suggest that Kidins220 is specifically necessary for *λLC* locus opening and/or its transcription in developing B cells. Mechanistically, this regulation is independent of the transcriptional regulation of elements of the recombination machinery.

### Subhead 3: Kidins220 is required for the generation of λLC^+^ B cells in κ-KO mice

We next aimed to force the development of λLC^+^ B cells in a Kidins220-deficient background by crossing CTRL and B-KO mice to κ-deficient mice. We used the well described iEκT model, where the intronic κ enhancer (iEκ) is replaced by a neomycin resistance cassette leading to a silenced *Igκ* locus (from now on indicated as κ-KO). Consequently, all developing B cells exclusively express the λLC (*62*). We used CTRL and B-KO mice heterozygous for the iEκT allele as littermate controls. We first analyzed the B cell compartment in the BM of all four genotypes (**Fig. 3A**). As shown in **Figure 3B-G**, the heterozygous ablation of one of the *κ* alleles did not overall change the already described phenotype of CTRL and B-KO mice (*37*). B-KO mice showed B cell frequencies and cell numbers comparable to CTRL but a severe reduction of λLC^+^ B cells (**Fig. 3C** and **G; Fig. S3A, B, G**). In both CTRL and B-KO mice, κ-KO did not significantly alter the total amount of B220^+^ B cells in the BM (**Fig. S3A, B**). However, a severe developmental block was observed in the generation of BCR^+^ B cells in both κ-KO genotypes shown by a decreased proportion of IgM^+^ (immature and mature) B cells and a corresponding increase in pro-/pre-B cells (**Fig. 3D**). The total pro-/pre-B cell numbers remained unchanged (**Fig. S3C**). Remarkably, the reduction in IgM^+^ cells in κ-KO mice was stronger in B-KO mice, representing only 15% of the B220^+^IgM^+^ population of the CTRL κ-KO mice (**Fig. 3E; Fig. S3D, E**). As expected, neither CTRL nor B-KO mice generated κLC-bearing B cells in the κ-KO background (**Fig. 3F; Fig. S3F**). Instead, all generated BCR^+^B cells used the λLC (**Fig. 3B** and **G**). Indeed, this data explain the remarkable loss of IgM^+^cells in B-KO κ-KO double KO mice (**Fig. 3E**). As previously described, κ-KO amplified λLC usage in CTRL mice (*62*). Although some amplification also occurred in Kidins220-deficient B cells, the amount of λLC^+^ B cells was severely reduced compared to CTRL (**Fig. 3G; Fig. S3G**).

**Fig. 3.**
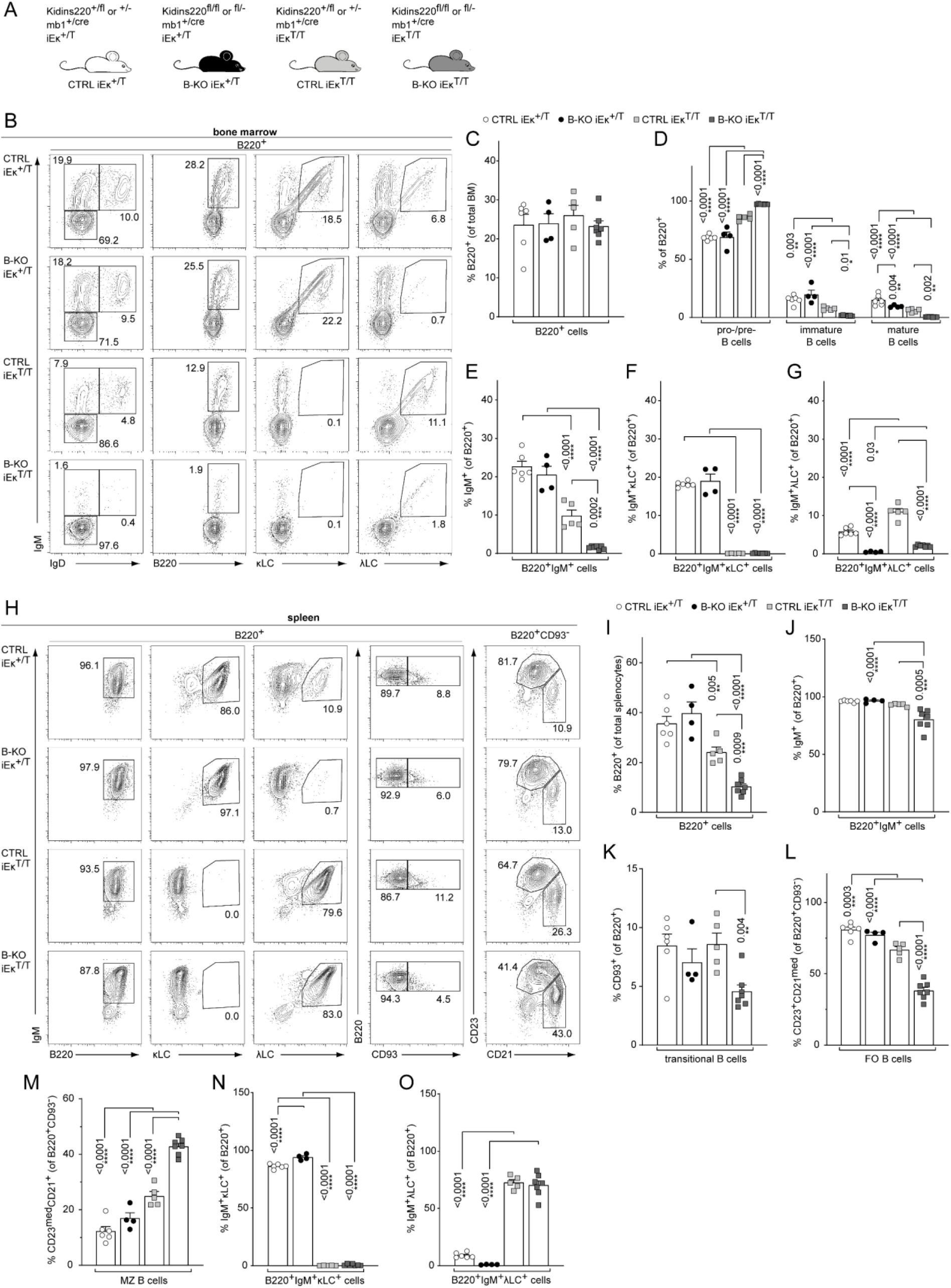
Kidins220 is required for the generation of λLC-bearing B cells even in κ-KO mice. (**A**) Genotypes of the *Kidins220* locus, the *κLC* locus and the *mb1* locus (cre-recombinase transgene) for CTRL and B-KO mice on heterozygous (iEκ^+/T^) or homozygous (iEκ^T/T^) κ-KO backgrounds. Representative flow cytometric analysis of the BM (**B**) and spleen (**H**) of CTLR and B-KO mice in κ-sufficient (iEκ^+/T^) and -deficient (iEκ^T/T^) backgrounds. (**C-G**; **I-O**) Pro-/pre-B cells were defined as B220^+^IgM^-^IgD^-^, immature B cells as B220^+^IgM^+^IgD^-^, mature B cells as B220^+^IgM^+^IgD^+^, transitional B cells as B220^+^CD93^+^, follicular (FO) B cells as B220^+^CD93^-^IgM^+^CD21^med^CD23^high^ and marginal zone (MZ) B cells as B220^+^CD93^-^IgM^+^CD21^high^CD23^med^. Quantification of three independent experiments with n = 4-7 mice per genotype. In all graphs, the mean ± SEM is plotted. Statistical analysis was performed using ANOVA test. CTRL and B-KO in iEκ^+/T^ or iEκ^T/T^ mice were compared. iEκ^+/T^ and iEκ^T/T^ mice were compared for CTRL or B-KO backgrounds. Only P values < 0.05 are indicated.

Immature IgM^+^ B cells emerge from the BM and become transitional B cells, which further mature in the spleen. Although the total splenocytes number did not significantly change between κ-sufficient and -deficient backgrounds in CTRL mice, κ-KO mice showed an overall reduction of the B220^+^ and B220^+^IgM^+^ B cell percentages and numbers in the spleen (**Fig. 3H-J**; **Fig. S3H-J**), concomitant with a relative increase in T cells (data not shown) (*62*). This reduction in B220^+^ B cells was strongly amplified by Kidins220-deficiency (60-80% reduction of B220^+^ B cells in B-KO κ-KO compared to CTRL κ-KO) (**Fig. 3I**; **Fig. S3I**). Silencing of the *κLC* locus, or deletion of Kidins220 expression in B cells, diminished transitional B cell numbers but not their percentages (**Fig. 3K**; **Fig. S3K**). The double KO enhanced these effects on transitional B cells (**Fig. 3K**; **Fig. S3K**). In the spleen, most of the transitional B cells further mature into follicular (FO) B cells, whereas a smaller fraction develops into marginal zone (MZ) B cells in the wild type situation (*63*). We previously reported that in the absence of Kidins220, maturation towards the MZ compartment is favored resulting in around 18% of all mature splenic B cells with a MZ B cell phenotype (**Fig. 3L** and **M**; **Fig. S3L, M**) (*37*). Here, we observed that κ-deficiency also favors the development of MZ B cells, and in κ-KO mice lacking Kidins220, this skewed differentiation toward the MZ compartment was further promoted (**Fig. 3M**; **Fig. S3M**). As expected, all B cells in the spleen exclusively express λLC BCRs in κ-KO mice, independent of Kidins220 expression (**Fig. 3N** and **O**; **Fig. S3N, O**). In the presence of Kidins220, development and maturation of λLC^+^ B cells allows for a compensation of B cell numbers reaching about 50% of the κLC-sufficient situation in most populations (**Fig. S3I-L**). However, in the absence of Kidins220, κ-KO mice contain significantly reduced λLC-expressing B cell numbers and remarkably failed to compensate to control levels (**Fig. S3O**). Taken together, Kidins220 is necessary in the BM for the development of λLC^+^ B cells. When the *κLC* locus cannot be rearranged, Kidins220 KO results in reduced B cell output from the BM.

### Subhead 4: BCL2 overexpression in Kidins220-deficient B cells partially rescues λLC usage *in vitro* and *in vivo*

For the generation of λLC^+^ B cells, the precursors should survive long enough to reach the state when the genes of the λLC becomes rearranged according to the generally accepted hypothesis that *κLC* gene rearrangement precedes *λLC* VJ recombination (*33*, *34*). To explore whether Kidins220-deficiency compromises the generation of λLC^+^ B cells by affecting B cell survival, we determined the amount of early apoptotic and dead cells throughout B cell development in the BM of CTRL and B-KO mice. As depicted in **Figure S4A**, Kidins220-deficiency showed a tendency towards increasing the amount of early apoptotic (Annexin V^+^PI^-^) and dead (Annexin V^+^PI^+^) cells during B cell development (except for pro-B cells), which became statistically significant for λLC^+^ B cells. This is in line with our previous report showing that *ex-vivo* cultured pro-/pre-B cells from B-KO mice have a decreased survival upon IL-7 withdrawal compared to CTRLs (*37*). Next, we aimed to revert the reduced survival of B-KO B cells by retrovirally transducing primary BM-derived pro-/pre-B cell cultures from CTRL and B-KO mice with a *BCL2-IRES-GFP* (or control *pMIG*) overexpression plasmid (**Fig. 4A**). Indeed, BCL2 overexpression rescued B cell survival from CTRL and B-KO mice to a comparable level over nine days following IL-7 withdrawal (**Fig. 4B**). In contrast, *pMIG* transduced cells showed a rapid reduction in viability, similar to untransduced primary cultures, whereby Kidins220-deficiency accelerated this process (**Fig. 4B**). Flow cytometric analysis of *BCL2*-transduced pro-/pre-B cell cultures revealed that IL-7 withdrawal induced κLC and λLC surface expression in both CTRL and B-KO B cells (**Fig. 4C**). However, even though *BCL2*-transduced B-KO B cells showed a pronounced increase of λLC^+^ B cells, it was still significantly reduced compared to CTRL cells. This reduction was compensated by a slightly enlarged κLC^+^ B cells proportion in B-KO B cells (**Fig. 4C**). We next analyzed the active states of the *LC* loci by determining the relative abundance of *κLC* and *λLC* germline transcripts. *BCL2*-transduced CTRL and B-KO B cells revealed a similar induction and kinetics of *κLC* germline transcripts (**Fig. 4D**). In contrast, Kidins220-deficiency dampened the abundance of *λLC* germline transcripts compared to CTRLs, even though the relative abundance of *λLC* germline transcripts seemed to be partially rescued after nine days of IL-7 withdrawal (**Fig. 4D**). Again, these effects were not caused by an altered gene expression of proteins and transcription factors of the recombination machinery such as *E12*, *E47*,*Rag1* and *Rag2* (**Fig. S4B**). Together, these data confirm a role for Kidins220 in the survival of B cell precursors. However, the function of Kidins220 seems to be beyond just facilitating B cell survival, since BCL2 overexpression did not fully rescue λLC expression in Kidins220-deficient primary B cell cultures.

**Fig. 4.**
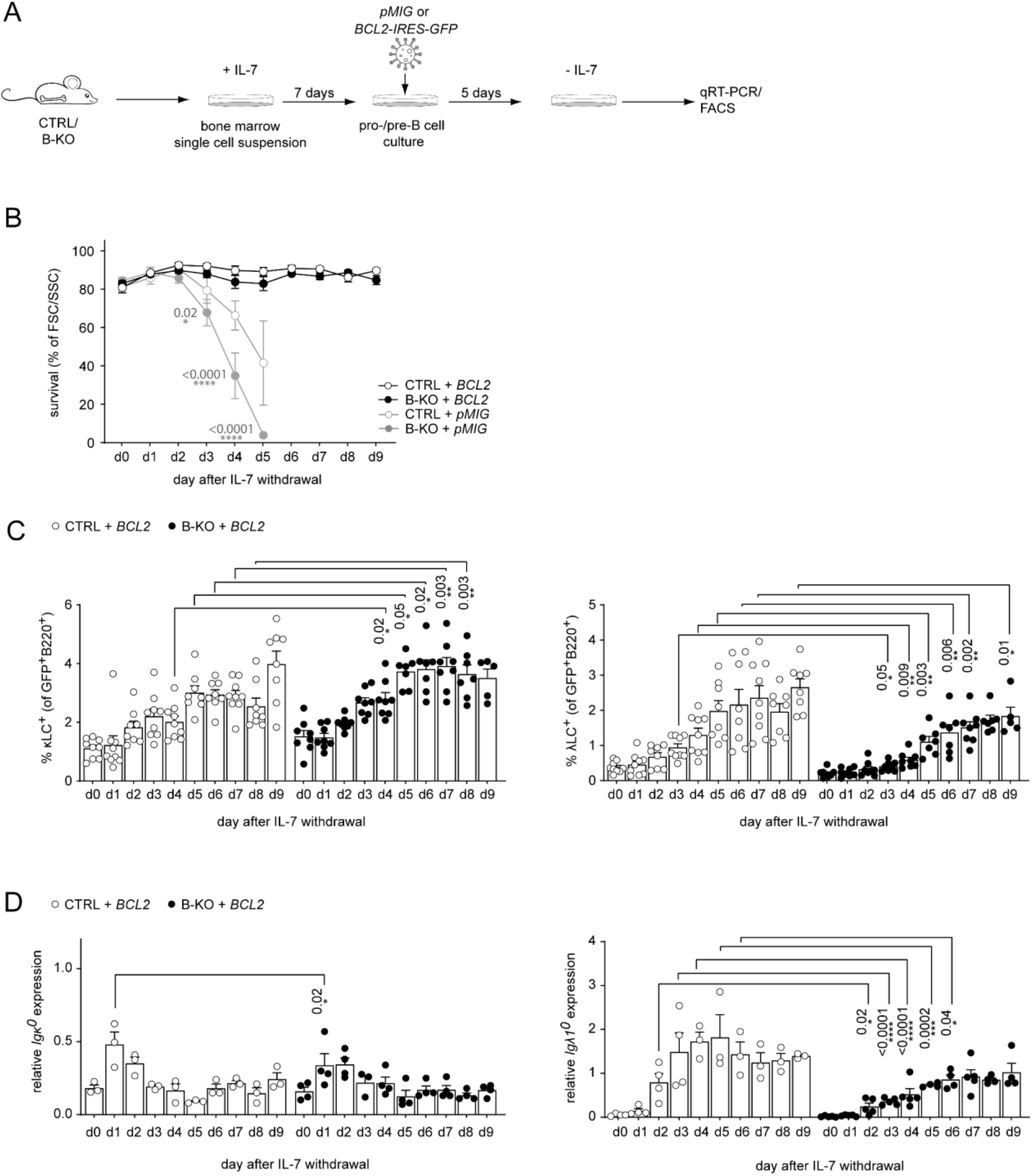
BCL2-mediated survival partially rescues λLC deficiency *in vitro*. (**A**) Experimental setup to generate primary BM-derived pro-/pre-B cell cultures overexpressing *BCL2*. Total BM was isolated from CTRL and B-KO mice and cultured for seven days in medium supplemented with IL-7 as in Figure 2. The cells were retrovirally transduced with a *BCL2-IRES-GFP* or *pMIG* overexpression plasmids and cultured in the presence of IL-7 for five more days. Upon IL-7 withdrawal, samples were collected directly (day 0) and then every 24 hours for nine days for analysis. (**B**) The percentage of living cells was estimated using flow cytometry based on FSC/SSC daily (three to five independent experiments; n = 5-11 mice per genotype). (**C**) Surface expression of λLC and κLC in *BCL2*-transduced pro-/pre-B cell cultures assayed by flow cytometry (four independent experiments; n = 5-9 per genotype; each symbol represents one mouse). (**D**) RNA was isolated, and reverse transcribed to quantify the relative expression of germline transcripts of *Igκ* and *Igλ* by qRT-PCR. Data were normalized to *Hrpt* (n = 3-4; each symbol represents one mouse. In all graphs, the mean ± SEM is plotted. Statistical analysis compared CTRL and B-KO samples for each day using ANOVA test. Only *P* values < 0.05 are indicated.

Next, we crossed the CTRL and B-KO mice to *BCL2*-transgenic mice, in which the *BCL2*-transgene flanked by the *vav* promoter results in BCL2 expression in all nucleated cells of hematopoietic origin (*64*) (**Fig. 5A**). BCL2 overexpression increased the percentage and total number of B cells in the BM in CTRL mice (**Fig. 5B** and **C**; **Fig. S5A, B**) as previously reported (*64*). However, this was not the case in B-KO mice (**Fig. 5B** and **C**; **Fig. S5A, B**). IgM^+^ mature B cells clearly accumulated in the BM of CTRL *BCL2*-transgenic mice as previously shown (*65*, *66*) (**Fig. 5D** and **E**; **Fig. S5C, D**). The mature IgM^+^ B cell population in the BM of B-KO *BCL2*-transgenic mice showed a similar tendency towards expansion, although total cell numbers remained significantly lower than in CTRL mice (**Fig. 5D** and **E**; **Fig. S5C, D**). Further characterization of the IgM-expressing B cells revealed that the percentages and total cell numbers of both κLC^+^ and λLC^+^ B cells were expanded in *BCL2*-transgenic CTRL as described (*27*, *36*) (**Fig. 5F** and **G; Fig. S5E, F**). In B-KO mice, *BCL2*-transgenic expression also increased the percentage and cell numbers of κLC^+^ and λLC^+^ B cells (**Fig. 5F** and **G** and **Fig. S5E, F**). However, that did not reach the levels of κLC^+^ B cell numbers, and of the percentage and total number of λLC^+^ B cells observed in the CTRL *BCL2*-transgenic mice (**Fig. 5F** and **G** and **Fig. S5E, F**). In line with a previous report (*35*), the expansion of the λLC^+^ compartment in both genotypes indicated a strong dependency of λLC^+^ B cells on B cell survival (expansion of total cell numbers due to BCL2 overexpression between 6-14-fold for λLC^+^ B cells compared to 2-4-fold for κLC^+^ B cells) (**Fig. S5G**). Furthermore, the increased fold change of the relative numbers of λLC^+^ B cells in B-KO compared to CTRL mice due to BCL2 overexpression suggests a role of Kidins220 in promoting B cell survival. The BCL2 overexpression-driven increase in B cell precursors in the BM of CTRL mice resulted in the increased total number of splenocytes, IgM^+^, κLC^+^ and λLC^+^ B cells as reported (*64*–*66*) (**Fig. S5H-L**). Ectopic BCL2 expression in the B-KO background did not change the absolute counts of B cell subpopulations, except for a reduction in MZ B cells (**Fig. S5H-O**). The percentage of B cells in the spleen was slightly reduced in both CTRL and B-KO mice because of a concomitant expansion of T cells (**Fig. 5H** and **I**; data not shown). In the four genotypes analyzed, splenic B cells were IgM positive (**Fig. 5J**). The amount of transitional B cells was increased in BCL2-overexpressing CTRL mice (**Fig. S5M)**, but their proportion within the B220^+^ B cells was reduced both in CTRL and in Kidins220-deficient mice compared to non-transgenic mice (**Fig. 5K**; **Fig. S5M**). Furthermore, enforced BCL2 expression severely reduced the MZ B cell compartment in CTRL mice and B-KO mice (**Fig. 5L** and **M**; **Fig. S5N, O**) (*67*, *68*). Most importantly, BCL2-overexpression led to a relative decrease of κLC^+^ B cells accompanied with a relative increase of λLC^+^ B cells in both CTRL and B-KO mice (**Fig. 5N** and **O**; **Fig. S5P**). For the CTRLs, this is in line with previous reports in other *BCL2*-transgenic mouse models (*27*, *69*).

**Fig. 5.**
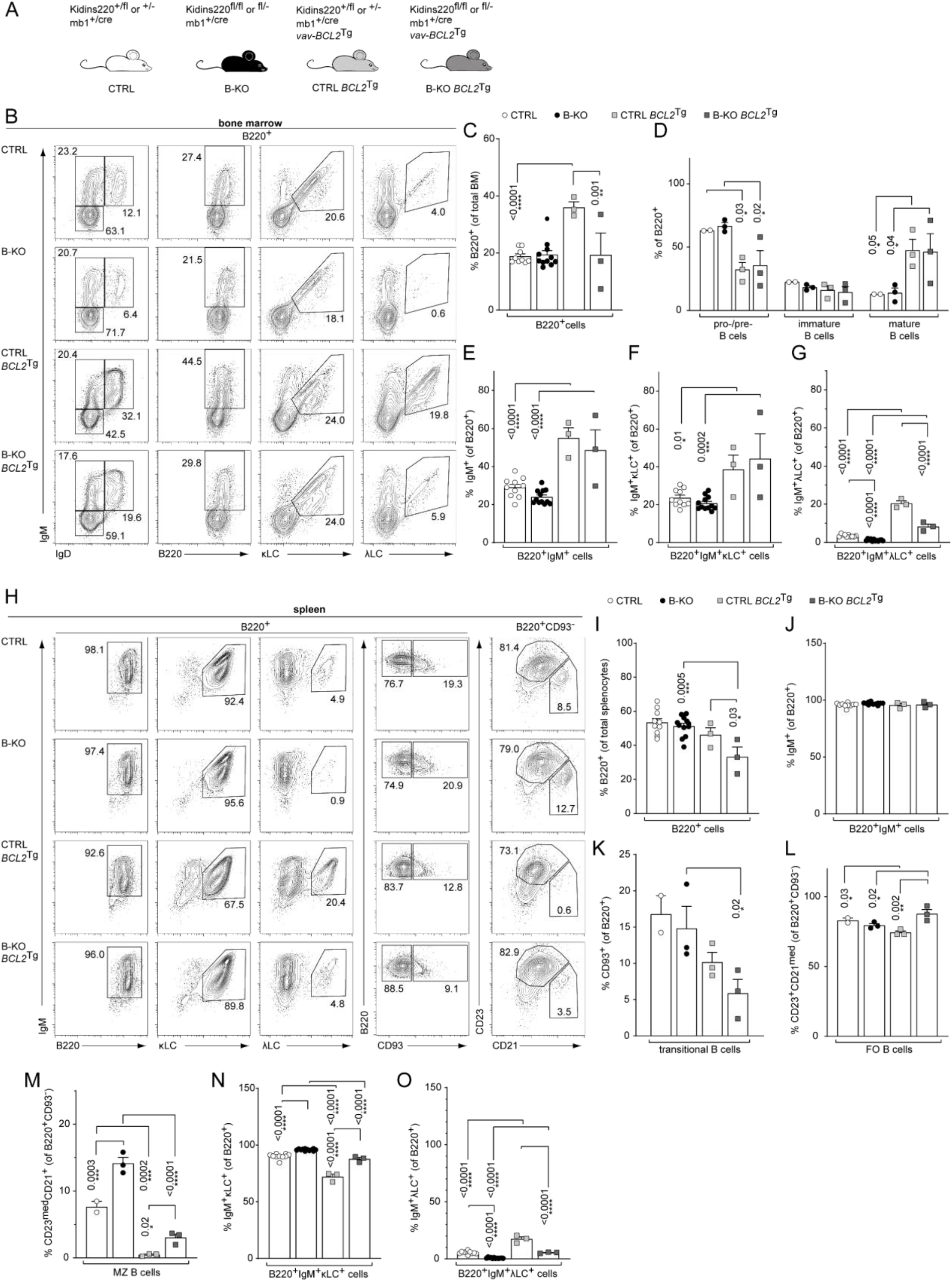
Ectopic BCL2-expression partially rescues λLC expression in Kidins220 B-KO mice. (**A**) The genotypes of the *Kidins220* locus, the *mb1* locus (cre-recombinase transgene) and *vav-BCL2* transgene (vav-BCL2^Tg^) expression for CTRL and B-KO mice are depicted. Representative flow cytometry plots of CTRL and B-KO mice in the absence or presence of the BCL2 transgene of BM (**B**) and spleen (**H**). (**C-G**; **I-O**) Pro-/pre-B cells are defined as B220^+^IgM^-^IgD^-^, immature B cells as B220^+^IgM^+^IgD^-^, mature B cells as B220^+^IgM^+^IgD^+^, transitional B cells as B220^+^CD93^+^, follicular (FO) B cells as B220^+^CD93^-^IgM^+^CD21^med^CD23^high^ and marginal zone (MZ) B cells as B220^+^CD93^-^IgM^+^CD21^high^CD23^med^. Quantification of three to five independent experiments with n = 2-12 mice per genotype; each dot represents a mouse. In all graphs, the mean ± SEM is plotted. Statistical analysis was performed using ANOVA test. CTRL and B-KO in the presence or absence of the BCL2 transgene were compared. Non-transgenic and BCL2^Tg^ mice were compared for CTRL or B-KO backgrounds. Only P values < 0.05 are indicated.

Taken together, enforced survival by overexpression of the anti-apoptotic protein BCL2 enhanced the generation of λLC^+^ B cells in CTRL and B-KO mice. The increased proportion of λLC^+^ B cells in Kidins220-deficient B cells overexpressing BCL2, did not restore total numbers of peripheral λLC^+^ B cells to control levels implying a role for Kidins220 in the generation of λLC^+^ B cells beyond just regulating B cell survival.

### Subhead 5: Kidins220 facilitates λLC expression during *in vivo* receptor editing

The amount of λLC^+^ B cells in the repertoire is also determined by the extent of receptor editing during the establishment of tolerance (*33*). Autoreactive BCRs transmit signals upon self-antigen recognition resulting in the elimination of the autoreactive BCR from the repertoire by replacing the LC with an innocuous one by secondary VJ-gene rearrangement at the *κLC* or *λLC* locus. To analyze the potential role of Kidins220 during tolerance induction *in vivo*, we used a previously described model characterized by the ubiquitous transgenic expression of a membrane bound anti-Igκ-reactive single chain antibody: the κ-macroself mice (*35*). By using κ-macroself mice as recipients for BM reconstitutions from either CTRL or B-KO mice, all developing donor B cells expressing a κLC will recognize the ubiquitously expressed κ-transgene as a self-antigen, initiating receptor editing and generating B cells with solely λLC surface expression to maintain immune tolerance (*35*). We isolated hematopoietic stem cells (HSC) from CTRL and B-KO mice (both expressing the leukocyte marker CD45.2) and injected them into sublethally irradiated CD45.1^+^ wild type (WT) or CD45.1^+^κ-macroself mice (**Fig. 6A**). The HSC of CTRL and B-KO mice gave rise to similar percentages of B220^+^ B cells in the BM of both CD45.1^+^ WT and κ-macroself transgenic mice (**Fig. 6B** and **C**), with a slight increase in total B cell numbers in κ-macroself mice (**Fig. S6A, B**). As expected, κLC^+^ B cells were rarely detected in κ-macroself mice reconstituted with either CTRL (*35*) or Kidins220-deficient HSC (**Fig. 6D**; **Fig. S6C**). Hence, recognition of strong self-antigen signals and successful elimination of autoreactive BCRs from the cell surface does not require Kidins220. The negative selection of κLC^+^ B cells was accompanied by a significant increase in λLC usage when CTRL HSC were transferred into κ-macroself mice (*21*, *35*) (**Fig. 6E**; **Fig. S6D**). Transfer of B-KO HSC to non-transgenic WT mice still showed the reduction in λLC-expressing B cells in the BM compared to CTRL mice (**Fig. 6E**; **Fig. S6D**). Thus, the diminished production of λLC^+^ cells in the absence of Kidins220 is B-cell intrinsic. In κ-macroself mice reconstituted with HSC from B-KO mice, total λLC^+^ B cell numbers were elevated compared to B-KO (**Fig. S6D**). However, the percentage and total number of λLC^+^B cells were still significantly decreased compared to their control counterparts (**Fig. 6E**; **Fig. S6D**). Interestingly, the percentage and total numbers of BCR^-^ B cells in the BM were significantly increased in κ-macroself mice independent of Kidins220 expression (**Fig. 6F**; **Fig. S6E**) as previously observed (*35*).

**Fig. 6.**
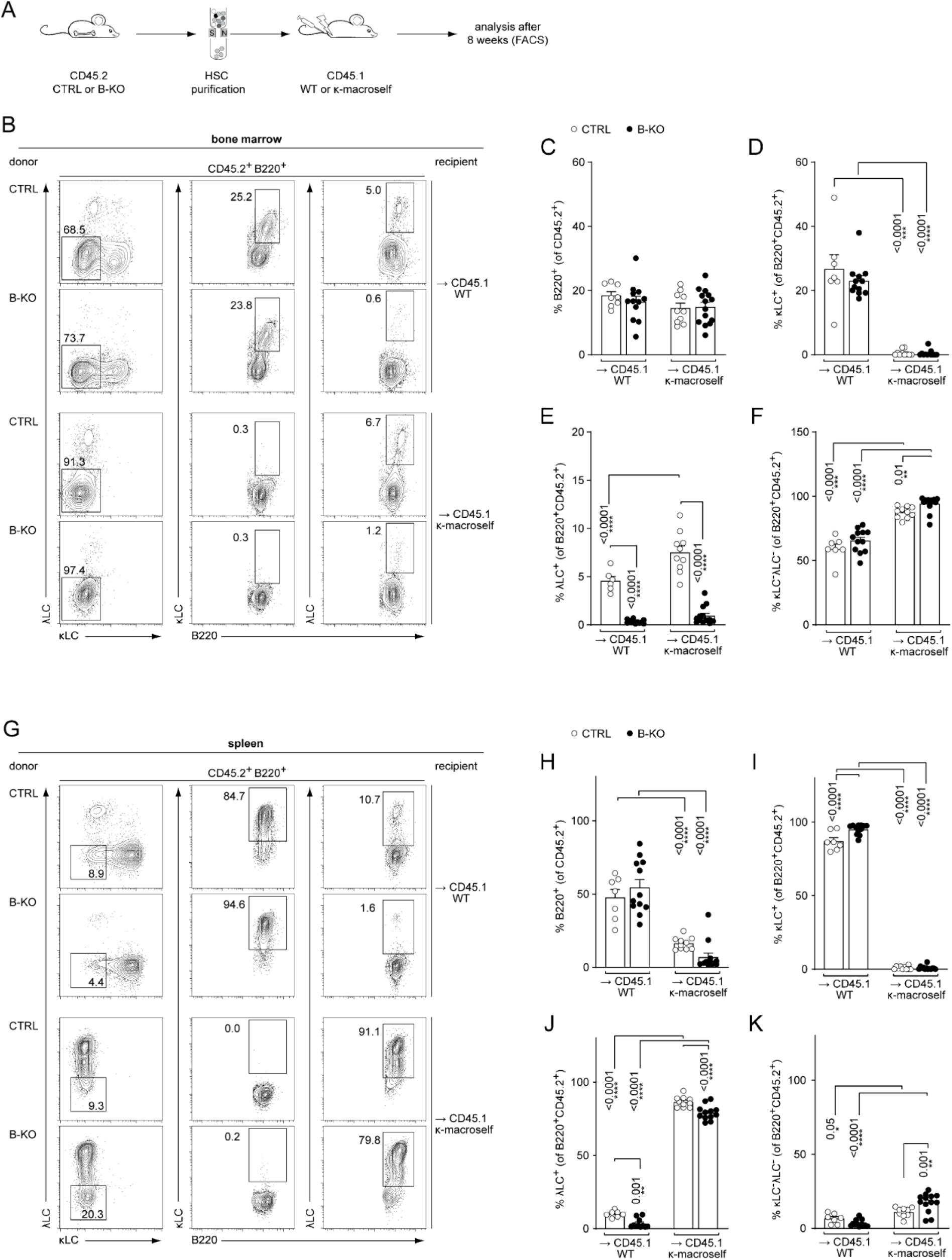
Kidins220 is dispensable for the elimination of autoreactive BCRs but necessary for the expression of innocuous λLC during tolerance induction. (**A**) Experimental setup for BM transfer into CD45.1 mice. HSC from CTRL and B-KO mice were isolated by negative magnetic purification. 5 × 10^5^ cells were injected intravenously into sublethally irradiated CD45.1 WT or CD45.1 κ-macroself transgenic mice. After eight weeks, the mice were sacrificed, and their organs were analyzed by flow cytometry. Representative plots of the BM (**B**) and spleen (**G**) are shown. (**C-F**; **H-K**) For the quantification, three independent experiments were pooled (n = 6-13 mice per condition). In all graphs the mean ± SEM is plotted. Statistical analysis was performed using ANOVA test. CTRL and B-KO in the presence or absence of the κ-macroself transgene were compared. WT and κ-macroself transgenic mice were compared for CTRL or B-KO backgrounds. Only *P* values < 0.05 are indicated.

The total B cell compartment in the spleens of these mice was reduced in percentage and numbers compared to non-transgenic littermates, most probably due to the reduced output from the BM (**Fig. 6G** and **H**; **Fig. S6F, G**) (*21*, *35*). The lack of κLC-expressing B cells in the periphery, independent of Kindins220 expression, indicated a functional tolerance induction (**Fig. 6I**; **Fig. S6H**). Concomitantly, there was an increase in λLC usage in both groups (**Fig. 6J**). However, a significant increase in λLC^+^ B cell numbers was observed only in κ-macroself mice reconstituted with HSC from CTRL mice (**Fig. S6I**). As in the BM, the relative number of splenic BCR^-^ B cells was increased, independent of Kidins220 expression in κ-macroself mice (**Fig.S. 6K**). This was reflected in total cell numbers for κ-macroself mice reconstituted with HSC from CTRL mice (**Fig. S6J**). These BCR^-^ B cells might have downregulated their BCR expression below the detection limit due to its recognition of the κ-macroself antigen. All in all, these data indicate that Kidins220 is not required for the transmission of signals downstream of the IgM-BCR upon recognition of strong self-antigens such as the chimeric anti-Igκ antibodies, and for the subsequent elimination of the autoreactive BCRs from the repertoire. However, Kidins220 seems to be essential for efficient expression of innocuous λLCs, possibly leaving “holes in the BCR repertoire”.

### Subhead 6: Ectopic BCL2 fails to rescue λLC expression during *in vivo* receptor editing

We next asked whether increased survival might rescue λLC expression during receptor editing. To this end, we modified our BM-transfer protocol by transducing the isolated HSC of CTRL and B-KO mice 24 hours post-isolation with a *BCL2-IRES-GFP* overexpression plasmid. Following another 24 hours of HSC culture in optimized medium, the cells were injected into sublethally irradiated CD45.1^+^ WT or κ-macroself mice for BM reconstitution (**Fig. 7A**). For the analysis, all cells were pre-gated on the GFP^+^ population (that marks BCL2-expressing cells) prior to gating for the individual B cell subpopulations. BCL2-overexpressing HSC from CTRL and B-KO mice reconstituted the BM of CD45.1^+^ WT and κ-macroself transgenic mice equally well, since we obtained similar B cell frequencies and total cell numbers in all conditions (**Fig. 7B** and **C; Fig. S7A**). As expected, ectopic BCL2 expression resulted in an increased relative number of both κLC^+^ as well as λLC^+^ B cells in CTRL and Kidins220-deficient B cells in the BM of non-transgenic WT mice when compared to non-transgenic WT mice reconstituted with non-BCL2-expressing HSC (**Fig. 7D** and **E**; **Fig. 6D** and **E**). Transgenic BCL2 expression in the presence of the anti-κ‘auto-antigen’ did not override the induction of central tolerance, in line with previous reports (*35*, *70*) (**Fig. 7D; Fig. S7B**). Instead, it led to a higher percentages and total cell numbers of λLC^+^ B cells in κ-macroself transgenic mice reconstituted with HSC from CTRLs (**Fig. 7E**; **Fig. S7C**). In contrast, neither the percentage nor the total number of λLC^+^ B cells in κ-macroself transgenic mice reconstituted with B-KO HSC increased upon BCL2-overexpression, nor did they reach the levels observed in mice reconstituted with HSC from CTRLs (**Fig. 7E**; **Fig. S7C**). The presence of the κ-macroself antigen only marginally increased the presence of BM B cells that lack surface expression of LC, and therefore BCR (**Fig. 7F**; **Fig. S7D**). All these findings from the BM were reproduced in the spleens with a similar phenotype (**Fig. 7G-K; Fig. S7E-H**).

**Fig. 7.**
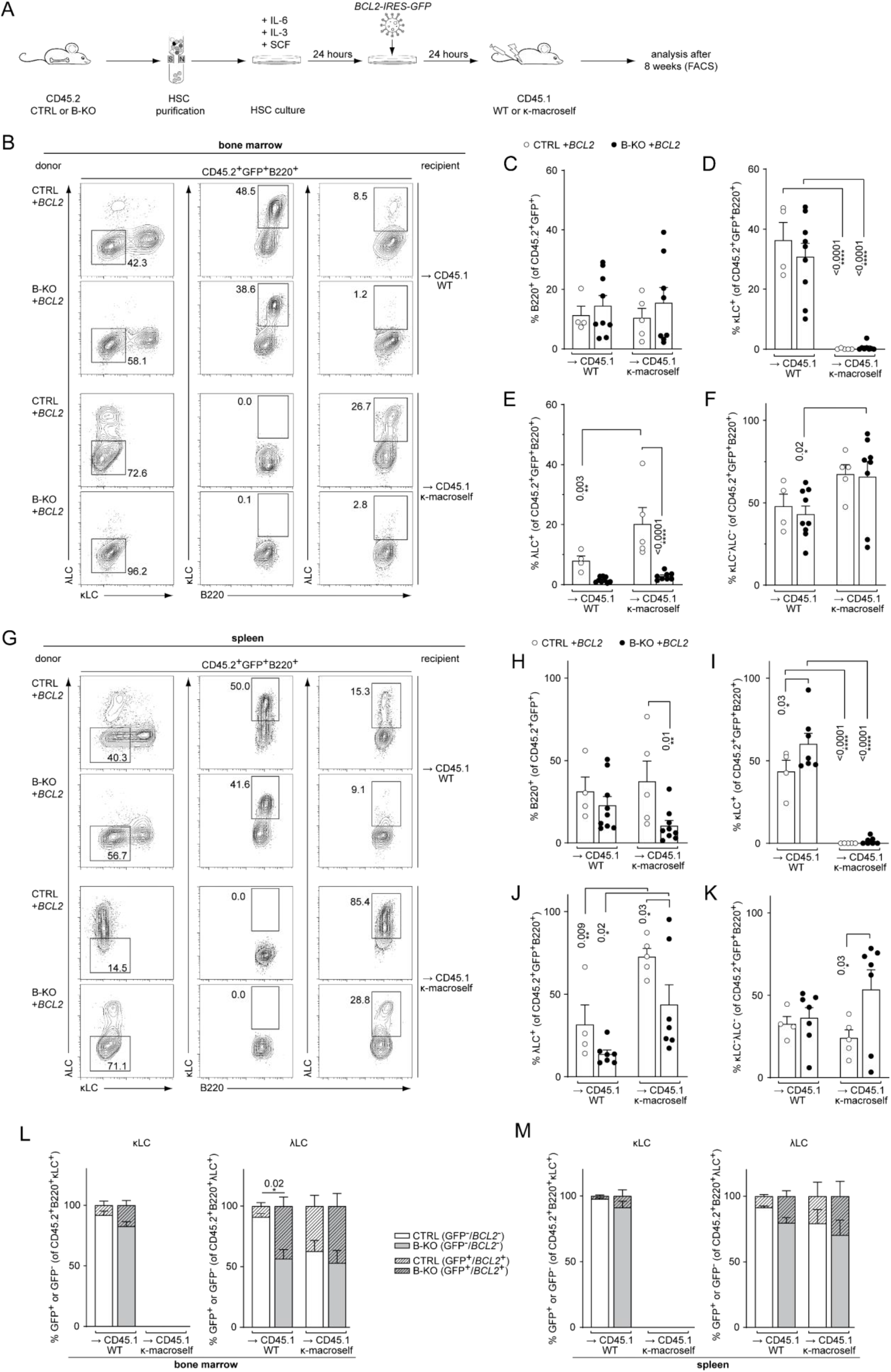
BCL2 overexpression fails to rescue λLC expression during tolerance induction. (**A**) Experimental setup for BM transfer of lentivirally transduced HSC into CD45.1 WT or CD45.1 κ-macroself transgenic mice. HSC from CTRL and B-KO mice were isolated by negative magnetic purification and cultured for 24 hours in the presence of IL-6, IL-3 and SCF. The cells were then lentivirally transduced with a *BCL2-IRES-GFP* overexpression plasmid and cultured for another 24 hours. 5 × 10^5^ cells were injected intravenously into sublethally irradiated CD45.1 WT or CD45.1 κ-macroself transgenic mice for BM reconstitution. After eight weeks, the mice were sacrificed, and their organs were analyzed by flow cytometry. (**B-K**) BCL2-expressing donor B cells were analyzed by pre-gating on GFP^+^ cells. Representative plots of BM (**B**) and spleen (**G**) and statistical analysis (**C-F**; **H-K**) are shown. (**L**), (**M**) Percent of donor B cells expressing BCL2 (GFP^+^) and not expressing BCL2 (GFP^-^) upon gating for κLC^+^ (left) or λLC^+^ (right) cells in the BM (L) and the spleen (M). Significance was determined by comparing the percentage of the GFP^+^ (striped) bars between the conditions. All panels are generated from two independent experiments (n = 4-9 mice per condition). In all graphs, the mean ± SEM is plotted. Statistical analysis was performed using ANOVA test. CTRL and B-KO in the presence or absence of the κ-macroself transgene were compared. WT and κ-macroself transgenic mice were compared for CTRL or B-KO backgrounds. Only *P* values < 0.05 are indicated.

We next investigated the contribution of prolonged survival in all four conditions by first gating on κLC^+^ or λLC^+^ B cells and subsequently gating on BCL2-overexpressing (GFP^+^; striped bars) *versus* non-overexpressing cells (GFP^-^; clear bars) (**Fig. 7L** and **M**). Kidins220-deficient λLC^+^B cells benefit more from the overexpression of BCL2 compared to CTRL counterparts in the BM of CD45.1^+^ WT mice. This difference however disappeared in the BM of κ-macroself transgenic mice (**Fig. 7L**). The same tendencies were observed in the spleens of the respective mice (**Fig. 7M**). These data revealed that BCL2 anti-apoptotic signaling has a greater impact on the generation of λLC^+^ B cells than of B cells carrying a κLC BCR. Taken together, these data support a model in which Kidins220 is dispensable for the transmission of strong BCR-mediated auto-antigenic signals during tolerance induction. However, Kidins220 is needed to rearrange the genes of the *Igλ* locus. Even though Kidins220-deficient B cells have a reduced life-span, prolonged survival by transgenic BCL2 overexpression alone, or in combination with enforced receptor editing, failed to fully rescue λLC-expression.

### Subhead 7: Kidins220 is required for optimal pre-BCR signaling

*LC* locus opening and successful recombination depends on optimal pre-BCR and/or BCR signaling. We previously demonstrated that Kidins220 is important for BCR-mediated downstream signaling of *ex vivo* stimulated primary splenic B cells (*37*). Further, mice deficient for pre-BCR downstream signaling components, such as BTK, SPL65 or PLCγ2, show an impaired upregulation of surface markers like CD25, and major histocompatibility complex (MHC) class II concomitant with defective downregulation of CD43 (*16*, *26*, *29*, *57*, *71*). Thus, we analyzed the surface expression of these markers on BM B cells from CTRL and B-KO mice. Indeed, developing B cells of B-KO mice showed a reduced percentage of CD25^+^ cells with slightly lower levels of CD25 on the B cell surface (**Fig. 8A**). Furthermore, B-KO mice failed to efficiently upregulate MHCII, whereas the relative amount of CD43-expressing B cells was increased (**Fig. 8A** and (*37*)). This indicates reduced pre-BCR signaling. To further support these findings, we analyzed the basal level of reactive oxygen species (ROS) in CTRL and B-KO mice. ROS serve as important second messengers (reviewed in (*72*)). To this end, we isolated primary BM cells and splenocytes from B-KO and CTRL mice and labeled them with the fluorogenic dye DCFDA, in combination with a surface marker to analyze cellular ROS production within the different stages of B cell development. B cells from B-KO mice showed a significantly reduced level of ROS during almost all BM developmental stages except for large pre-B cells (**Fig. 8B**). We confirmed a significant drop of ROS from the large to small pre-B cells, as previously reported (*73*). This drop was much more pronounced in Kidins220-deficient developing B cells. ROS were maintained at very low levels throughout B cell development in the BM of Kidins220-deficient mice. Splenic naïve B cells of Kidins220-sufficient and -deficient mice showed similar low ROS levels. These results might indicate that Kidins220-deficient B cells produce less ROS due to impaired pre-BCR and BCR-mediated signaling. Reduced pre-BCR and BCR signaling might in turn prevent the opening and recombination of the *λLC* locus in developing B cells.

**Figure 8:**
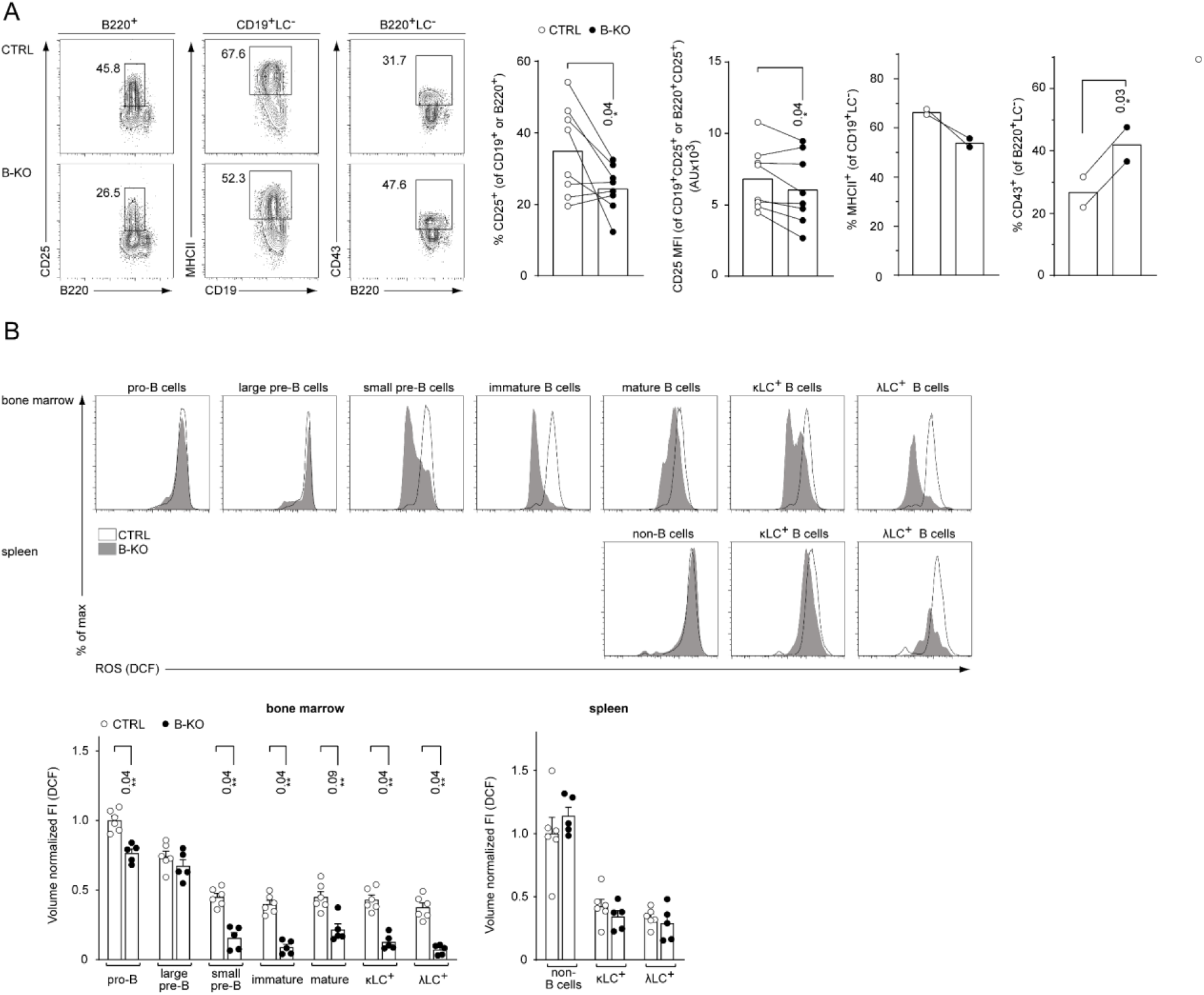
Reduced pre-BCR signaling in the absence of Kidins220. (**A**) Representative flow cytometry plots of the BM of CTRL and B-KO mice showing the surface expression of CD25, MHCII and CD43 (left). For CD25, quantification of eight independent experiments with n = 12-14 mice per genotype is shown. Each dot represents the average of an individual experiment. For MHCII and CD43, quantification of two independent experiments with n = 2 mice per genotype is shown. Each dot represents an individual mouse. Statistical analysis was performed using Wilcoxon matched-pairs signed rank test or paired Student’s *t*-test test comparing CTRL and B-KO mice (**B**) B cell subpopulations in the BM and spleens of CTRL and B-KO mice were analyzed by flow cytometry using specific antibodies against B220, CD117 (c-kit), CD25, κLC and λLC. Cells were additionally stained with DCFDA to assess ROS levels. Representative histograms are shown on the top. For quantification (bottom), fluorescent intensities of the indicated metabolic marker were first normalized to the mean cell volume of each subpopulation and then normalized to pro-B cells (BM) or non-B cells (spleen). Three independent experiments were pooled; n = 5-6 mice per genotype. Statistical analysis was performed using Mann-Whitney test comparing CTRL and B-KO mice for a given cell population. In all graphs, the mean ± SEM is plotted. Only *P* values < 0.05 are indicated.

## DISCUSSION

We have previously discovered that the transmembrane protein Kidins220 binds to the BCR and regulates BCR signaling (*37*). Deleting Kidins220 in B cells (B-KO mice) results in a severe reduction of λLC B cells despite normal generation of κLC B cells, generating thus a skewed antibody repertoire. Our data indicate that the absence of Kidins220 specifically prevents opening and/or transcription of the *λLC* locus. Remarkably, Kidins220-deficient B cells fail to open and recombine the *λLC* locus, even in genetic scenarios where the *κLC* genes cannot be rearranged (κ-KO). Our single cell analysis of the frequencies of *Igλ* V-gene recombination has confirmed a clear preference for the recombination of *V_λ_1*-gene in WT mice (*48*, *49*). This also holds true for B-KO mice. The current model proposes that this preference results from the cooperative function of the λ_1-3_ and λ_2-4_ enhancers, which surround the *Igλ* V-gene segments 3’ and 5’ of the λ1 and λ3 family and are bound by the same transcription factors (*74*, *75*). The distance of the λ_2-4_ enhancer to the λ_1-3_ enhancer might preclude that *λLC* gene recombination 5’ of the λ_2-4_ enhancer profit from transcription factors and co-factors of the recombination machinery bound to the λ1-3 enhancer. The rearrangement of the *V_λ_1* to *JC_λ_1*-gene segment eliminates the intervening sequence comprising the *JC_λ_3*-gene segment, resulting in the accumulation of λ1LC B cells. Interestingly, we observed a relative increase in the usage of the genes of the λ3-family compared to the λ2-family in B-KO *versus* CTRL mice. This observation suggests that the recombination of the *Igλ* V-gene segments that are 5’ of the λ_2-4_ enhancer are more depended on Kidins220 expression than the others. A possible explanation for this observation is that pre-BCR and BCR signaling are reduced in the absence of Kidins220 making the recombination of λ2-family V*-* genes unlikely. Instead, *V_λ_1* is recombined to *JC_λ_3* more often in Kidins220-deficient mice. Together, our single cell sequencing analysis supports that Kidins220 is dispensable for *κLC*, but essential for efficient *λLC* locus opening and gene recombination.

Reduced production of λLC B cells has been previously described in studies using mouse models with genetic deletion of signaling molecules downstream of the pre-BCR and BCR like BTK, SLP65 and PLCγ2 (*15*, *16*, *26*–*29*, *57*). In contrast, the combined deletion of some of these signaling components almost completely abolishes both *λLC* and *κLC* germline transcription, V-gene recombination, and protein expression (*16*, *26*, *29*). These data led to the interpretation that the generation of λLC B cells is more sensitive to defects in the signals transmitted by the pre-BCR and/or BCR (*16*). Kidins220-deficiency has been previously described to dampen signaling by receptors in neurons, glial cells, adipocytes, T and B cells (*37*, *40*, *76*–*80*). Indeed, BCR stimulation in a Kidins220 knockdown mature B cell line resulted in dampened activation of the RAS-ERK pathway due to an inefficient coupling of the BCR to the downstream Raf kinases B-Raf and Raf-1 (*37*). Likewise, IgM-BCR-mediated stimulation of primary splenocytes from B-KO mice resulted in less phosphorylation of ERK and PLCγ2, as well as reduced calcium influx compared to CTRL cells; further highlighting the role of Kidins220 in BCR signaling (*37*). Our data herein suggest that pre-BCR signaling is also reduced in the absence of Kidins220. On the one hand, B-KO mice showed a reduction of the amount of MHCII^+^ and CD25^+^ cells as well as reduced CD25 surface expression on B cells in the BM. CD25 is upregulated upon pre-BCR signaling (*81*, *82*), and its expression level is reduced in mice with genetic ablations of BTK, SLP65, or PLCγ2 (*16*, *26*, *29*, *57*, *71*). On the other hand, Kidins220-deficient B cells showed reduced ROS levels during BM development, especially in those stages where pre-BCR (small pre-B cells) and IgM-BCR signaling (immature B cells) is crucial. ROS are important second messengers (reviewed in (*72*)). In mature B cells, BCR activation induces ROS production to fully activate NF-κB signaling (*83*, *84*). Considering the parallels between BCR and pre-BCR signaling, ROS might exert a similar role downstream of the pre-BCR. Indeed, previous reports have demonstrated that λLC B cell development but not κLC editing depends on NF-κB signals in line with our results (*36*). Taken together, we hypothesize that Kidins220 mediates its effects by enhancing pre-BCR and BCR signaling although, we cannot formally rule out that Kidins220 might possess direct, signaling-dependent or -independent effects on the *λLC* locus opening.

In addition, our data suggest that Kidins220 contributes to pre-B cell survival, which is necessary for *IgLC* gene recombination. In fact, a prolonged life span in pre-B cells, induced by overexpression of anti-apoptotic BCL2, increases the amount of λLC^+^ B cells (*27*, *35*, *36*). BCL2 overexpression rescues the generation of λLC B cells in scenarios in which NF-κB signaling was abolished during B cell development (*36*). A connection between pre-BCR and BCR signaling and survival is well recognized (*85*–*87*). However, there is limited data on the exact mechanism. In particular, pre-BCR signaling might lead to the activation of the NF-κB pathway, that in turn regulates the transcription of *Pim2*, a pro-survival protein in pre-B cells (*36*, *88*, *89*). Pre-BCR-mediated ERK activation might induce phosphorylation of the pro-apoptotic protein BIM and thereby interfere with its inhibitory interaction with BCL2, resulting in enhanced pre-B cell survival similar to reports on IgM-BCR signaling (*1*, *90*, *91*). In addition, pre-BCR-mediated ERK signaling might induce *Bcl2* transcription itself, thereby prolonging B cell survival during B cell development (*87*, *91*). Thus, the afore-discussed defects in pre-BCR and/or BCR signaling in Kidins220-deficient B cells could lead to BCL2 levels that are insufficient to protect developing B cells from cell death. Indeed, *ex vivo* cultured B cell precursor from B-KO mice die faster upon IL-7 withdrawal (*37*), and we also detected increased amounts of apoptotic cells during BM B cell development in these mice. However, overexpression of BCL2 did not fully rescue λLC-deficiency in B-KO mice, suggesting that prolong survival of B cell precursors is required, but not sufficient, to optimally generate λLC B cells in the absence of Kidins220 expression.

The κLC to λLC ratio is also affected by receptor editing. Receptor editing can lead to further recombination on either the same *κLC* allele, the second *κLC* allele or can proceed to the *λLC* locus (*34*). The κ and λLCs differ in the physicochemical and structural properties of their CDR3 regions, and λLCs are proposed to be more effective in rescuing BCRs that show autoreactivity (*92*, *93*). In line with previous studies, we show here that genetically forcing receptor editing using κ-deleting (κ-macroself) mice increased the production of B cells using the λLC in Kidins220 competent mice (*21*, *35*). In the absence of Kidins220, strong autoreactive ligands efficiently induce the downregulation of surface κLC^+^-BCRs indicating successful tolerance induction. However, the replacement of the autoreactive κLC by an innocuous λLC was severely impaired in the absence of Kidins220. These data underline the importance of Kidins220 for *λLC* gene recombination, even under the high selective pressure imposed by the κ-macroself mice. Indeed, overexpression of BCL2 did not profoundly enhance λLC expression by B-KO cells in the κ-macroself recipients, highlighting again that increasing survival is not sufficient to optimally obtain λLC B cells in the absence of Kidins220 expression. The data obtained in the κ-macroself mice are in line with our single cell repertoire analysis showing that we did not find any signs of potentially autoreactive BCRs in B-KO mice. Since Kidins220-deficient B cells do not open the *λLC* locus efficiently, secondary rearrangement at the *κLC* locus should be increased. Indeed, we observed a slight increase in the *J_κ_5-gene* usage in B-KO mice compared to CTRLs. *J_κ_5* is associated with several rounds of secondary rearrangement and often represents the last rearrangement before silencing of the *κLC* locus by recombining sequence recombination (*45*, *46*).

In conclusion, we suggest that Kidins220 might be key to support pre-BCR and BCR signaling to specifically regulate the generation of λLC B cells. It has been previously postulated that the opening of the *λLC* locus is more sensitive to a reduction in certain proteins such as IRF-4, E2A, and RAG1/2, whose activities are regulated by pre-BCR and BCR signals, than the *κLC* locus (*15*, *21*, *22*, *24*, *70*, *94*, *95*). In the absence of Kidins220, pre-BCR and BCR signaling might be reduced to a level that still allows efficient *Igκ-gene* rearrangement but fails to induce *Igλ-gene* rearrangement, after unsuccessful *κLC* gene rearrangement and/or receptor editing. Even though transcript levels of *Rag1/2*, *E12* and *E47* were similar between Kidins220-sufficient and -deficient B cells, the abundance of active RAG and/or E2A proteins might still be different. In fact, ERK phosphorylates and thereby activates E47 downstream of the IgM-BCR (*94*). Likewise, an active ERK pathway negatively regulates ID3 proteins, which in turn inhibit E47 activity (*11*). In line with this, it was shown that even though the transcription of E2A proteins was not changed in the absence of BTK, E2A binding to the iEκ was reduced in BTK-KO mice compared to littermate controls, leading to defects in *IgLC*-gene recombination (*16*). Thus, the afore-discussed defects in pre-BCR and BCR signaling in the absence of Kidins2020 might reduce the activation of RAG and/or E2A proteins, thereby dampening the *Igλ-gene* recombination.

To conclude, our study demonstrates that Kidins220 promotes pre-BCR and BCR signaling, supports pre-B cell survival, *λLC* locus opening and λLC expression during homeostatic B cell development and upon receptor editing. This study adds to our understanding of the underlying mechanism regulating the differential expression of the κ and λLCs in B cells and the generation of a self-tolerant repertoire.

## MATERIAL AND METHODS

### Cells and mice

Primary murine BM or pro-/pre-B cells were cultured in Opti-MEM™ containing 10% fetal calf serum (FCS), 2 mM L-Glutamine, 50 U/ml penicillin, 50 μg/ml streptomycin and 50 μM β-mercaptoethanol in a humidified saturated atmosphere at 37°C with 5% CO2. If needed, medium was supplemented with 5 ng/ml IL-7 (Peprotech). Primary murine HSC were cultured in complete DMEM GlutaMAX™ supplemented with 20% FCS, 50 U/ml penicillin, 50 μg/ml streptomycin, 50 μM β-mercaptoethanol and IL-3 (20 ng/ml, Peprotech), IL-6 (50 ng/ml, Peprotech) and SCF (50 ng/ml, Peprotech) in a humidified saturated atmosphere at 37°C with 7.5% CO2.

For lentiviral transduction, human embryonic kidney (HEK) 293T cells were used to produce retrovirus-containing supernatants. They were cultured in complete DMEM GlutaMAX™ medium supplemented with 10% FCS, 10 mM HEPES, 10 μM sodium pyruvate, 50 U/ml penicillin and 50 μg/ml streptomycin in a humidified saturated atmosphere at 37°C with 7.5% CO2.

The Kidins220mb1hCre (*37*), iEκT (*62*), vav-*BCL2*^Tg^ (*64*), C57BL/6-Ly5.1 (CD45.1 WT) and CD45.1 κ-macroself mice (*35*) were bred under specific pathogen-free conditions. All mice were backcrossed to C57BL/6 background for at least 10 generations. Mice were sex and age matched whenever possible. All animal protocols (G12/64) were performed according to German animal protection laws with permission from the responsible local authorities.

### Flow Cytometry and cell sorting

Single cells suspensions were gained from BM and spleens. Erythrocytes were removed prior to flow cytometry analysis by incubation in erythrocyte lysis buffer containing 150 mM NH4Cl and 10 mM KHCO_3_ for two (BM) or four (spleen) minutes at room temperature. 0.3 - 1 × 10^6^ cells were stained in PBS, containing 2% FCS and the respective antibodies for 20 min on ice. Cells were washed and measurements were performed using a Gallios (Beckman Coulter) or Attune NxT (Thermo Fisher Scientific) flow cytometer. Unspecific antibody binding was prevented by preincubation with TruStain FcX™ (clone 93, Biolegend).

Primary murine immature B cells for BCR repertoire analysis were sorted on a MoFlo Astrios EQ cell sorter (Beckman Coulter) using specific antibodies against B220, IgD and IgM (Fab Fragment). HSC were enriched by negative selection from total BM using biotinylated antibodies against CD3, B220, Ter119, CD11b, Ly6G/Ly6C followed by incubation with paramagnetic beads and magnetic cell sorting (MACS, Miltenyi Biotec).

### Antibodies

The following murine antibodies were used in flow cytometry: anti-B220(CD45R)-PECy7 (RA3-6B2) (eBioscience), anti-IgM-PE (eB121-15F9) (ThermoFisher Scientific), anti-IgD-eFluor450 (11-26c) (eBioscience), anti-IgM-Fab Fragment-Alexa Fluor 647 (Jackson ImmunoResearch Laboratories, Inc.), anti-Igλ1,λ2,λ3-FITC (R26-46) (BD Biosciences), anti-Igλ1,λ2,λ3-bio (R26-46) (BD Biosciences), anti-Igκ-V450 (187.1) (BD Biosciences), anti-CD117 (c-kit)-Brilliant Violet 421™ (2B8) (Biolegend), anti-CD25-APC (PC61.5) (ThermoFisher Scientific), anti-CD45.2-PE (104) (BD Biosciences), anti-CD45.1-APC (A20) (Biolegend), anti-CD3-biotin (145-2C11) (eBioscience), anti-B220(CD45R)-biotin (RA3-6B2) (eBioscience), anti-Ter119-biotin (TER-119) (eBioscience), anti-Ly6G/Ly6C (RB6-8C5) (eBioscience), anti-CD11b/Mac1 (M1/70) (eBioscience), anti-CD19-PB (6D5) (Biolegend), anti-MHCII-biotin (M5/114.15.2) (eBioscience), anti-CD43-PE (R2/60) (eBioscience). FITC Annexin V Apoptosis Detection Kit I was purchased from BD Biosciences and used according to manufacturer’s instructions.

### Primary BM cultures

Primary BM cultures were essentially generated by isolating total BM from the femur as previously described (*96*). Briefly, erythrocytes were removed by incubation in erythrocyte lysis buffer (see section Flow Cytometry) for 2 min at room temperature. 3 ml of 5 × 10^6^ cells/ml were cultured in one well of a p6 culture dish for seven days in the presence of 5 ng/ml IL-7 in complete Opti-MEM™. Fresh medium was added after 4 days. Afterwards, fresh medium was added every three days and cultures were split if needed. For IL-7 removal, cells were harvested and washed at least twice with an excess of medium without IL-7. Cells were plated in a p24 culture dish at a concentration of 2 × 10^6^ cells/ml in complete Opti-MEM™ for subsequent assays.

### Viral transduction

Murine retrovirus-containing supernatants were obtained by transfecting phoenix-eco cells using the PromoFectin (PromoKine) reagent according to manufacturers protocol using pMIG-hBCL2-IRES-GFP or control plasmids. Viral supernatant was collected and filtered after 48 hours and used directly. 1.5 × 10^6^ pro-/pre-B cells (or HSCs) were resuspended in 1 ml of viral supernatant containing Polybrene (1 μg/ml) and IL-7 (5 ng/ml) (or in complete HSC culture medium) and spin infected by centrifugation (90 min, 2500 rpm, 30°C). The viral supernatant was then removed, and cells were cultured under optimal conditions for subsequent assays or injection into mice 24 hours later.

Lentivirus was obtained by co-transfecting HEK 293T cells with LeGO-iG2-hBCL2-IRES-GFP, pCMVDR8.74 and pMD2G plasmids using Polyethyleneimine (PEI, Polysciences). Viral supernatant was harvested and combined 24- and 48 hours post-transfection. Lentiviral particles were enriched by overlaying in a 1:5 ratio on a 10% sucrose layer and centrifuging (4 hours, 10.000 x*g*, 8°C). The pellet containing lentiviral particles was resuspended in DMEM GlutaMAX™ w/o supplements and stored at −80°C. The viral titers were assessed by determining the multiplicity of infection (MOI). Briefly, 5 × 10^4^ HEK 293T cells per well were seeded in a p24 well plate in 1 ml medium. An aliquot of the concentrated virus was diluted 1:100 in medium. Various volumes (0, 1, 5, 10, 25 and 50 μl) of lentivirus dilution were added to the cells. After 48 h GFP expression was analyzed by flow cytometry and lentivirus titer was calculated using the following formula: Transduction units per ml = (number of cells x percent GFP^+^ cells x dilution factor) / (ml of lentivirus dilution). Primary HSC were spin infected with an MOI of 10 (90 min, 2500 rpm, 30°C) in complete HSC culture medium 24 hours prior to injection into the mice.

### qRT-PCR

Total RNA was isolated using TRIzol™ reagent (Thermo Fisher Scientific) according to manufacturer’s instructions. RNA concentration was assessed using Nanodrop. 1 μg of RNA was treated with DNase for 30 min at 37°C prior to cDNA synthesis. cDNA was prepared with oligo dT primers according to the manufacture’s protocol (Thermo Scientific). qRT-PCR was performed using Fast Start Universal SYBR Green Master (ROX) (Roche) according to manufacturer’s protocol. For amplification, gene-specific primers were used with a one-step protocol with an annealing temperature of 60°C. Expression levels were normalized to the expression of the house keeping gene *Hprt*.

### Primer

The following primers were used (in 5’ to 3’ direction): *Igκ^0^* for (CAGTGAGGAGGGTTTTTGTACAGCCAGACAG), *Igκ^0^* rev (CTCATTCCTGTTGAAGCTCTTGA), *Igλ1^0^* for (CTTGAGAATAAAATGCATGCAAG), *Igλ1^0^* rev (TGATGGCGAAGACTTGGGCTGG), *Rag1* for (ACCCGATGAAATTCAACACCC), *Rag1* rev (CTGGAACTACTGGAGACTGTTCT), *Rag2* for (ACACCAAACAATGAGCTTTCCG), *Rag2* rev (CCGTATCTGGGTTCAGGGAC), *E2A* for (GGGAGGAGAAAGAGGATGA), *E12* rev (GCTCCGCCTTCTGCTCTG), *E47* rev (CCGGTCCCTCAGGTCCTTC), *Hprt* for (GTTAAGCAGTACAGCCCCAAA), *Hprt* rev (AGGGCATATCCAACAACAAACTT).

### ROS staining

Single cell suspensions were isolated from BM and spleens. Erythrocytes were removed as described before (see section: Flow cytometry). 2 × 10^6^ cells were incubated in Opti-MEM with H2DCFDA (10 μM; Invitrogen™) for 30 min at 37°C. Afterwards, cells were washed twice and stained for surface markers for flow cytometry (see section: Flow cytometry). For quantification, the fluorescence intensity of H_2_DCFDA was normalized to the cell volume by FSC-W as an indicator of cell diameter as previously described (Stein *et al*, 2017; Tzur *et al*,2011).

### Data analysis

Flow cytometric data were analyzed using FlowJo V10 (Tree Star, Inc.) software. Data in Figures 1a and Figures 2–8 were statistically analyzed and displayed using GraphPad Prism9.

### Antibody repertoire library preparation and sequencing

For antibody repertoire analysis, the BM of three individual mice of each genotype (CTRL and B-KO respectively) were pooled and immature B cells (B220^+^, IgD^-^, IgM^+^) were FACS sorted using appropriate antibodies (or Fab-fragments for IgM). Single B cell antibody V(D)J libraries were prepared with the 10X Genomics Chromium Single Cell V(D)J platform version 1, allowing immune profiling of full-length antibody variable HC and LC (10XGenomics). Briefly, samples were loaded onto the Chromium Controller and partitioned into Gel Beads-in-emulsion (GEMs) containing single cells. The mRNA was reverse transcribed into barcoded cDNA. HC and LC full V(D)J variable sequences were amplified with a two-step PCR. Sequences were then fragmented and indexed with indices for Illumina sequencing. Quality control of the materials were obtained throughout the process and library concentration quantification was performed using an Agilent Bioanalyzer. Antibody library pools were sequenced on an Illumina MiSeq instrument at 2 × 300 bp paired-end reads using the MiSeq Reagent kit v3 (600 Cycles).

### Annotation, preprocessing and statistical analysis of antibody repertoires

Antibody repertoire data generated through the 10X Genomics V(D)J platform was demultiplexed using Cellranger mkfastq and subsequently annotated with IgBLAST version 1.14 and Cellranger version 4.0. Preprocessing included filtering for retaining CDR3s longer than 4 amino acids, selection of productive sequences and retaining only CDR3s occurring more than once in the repertoire, with productive sequences defined as sequences that are in-frame and contain no stop codons. Statistical analysis of antibody repertoire datasets was conducted using R 4.0.5. The Pearson’s correlation coefficients were calculated to measure the strength of the linear association between the *V*- and *J*-germline gene frequencies between CTRL and B-KO mice for HC and LC. The correlation is significant at the 0.05 level (2-tailed).

## Supplementary Materials

Fig. S1. Immature B-KO B cells show a skewed primary BCR repertoire.

Fig. S2. Kidins220 does not regulate E2A and Rag gene expression.

Fig. S3. Kidins220 is required for the generation of λLC bearing B cells, even in κ-KO mice.

Fig. S4. Kidins220 influences B cell survival.

Fig. S5. Ectopic BCL2 expression partially rescues λLC expression in B-KO mice.

Fig. S6. Kidins220 is dispensable for the elimination of autoreactive BCRs but is necessary for the expression of innocuous λLC during tolerance induction.

Fig. S7. BCL2 overexpression partially rescues λLC expression during tolerance induction.

## Acknowledgements

We thank K. Fehrenbach for technical assistance, the group of M. Erlacher, M. Reth and F. Cesca for providing mice and J. Jellusova for technical advice with ROS staining. We further want to thank core facilities at the BIOSS, CIBSS and ZTZ in Freiburg. We thank P. Nielsen, M. Reth, K. Schachtrup for stimulating discussions and intellectual input, and S. Pathan-Chhatbar for carefully reading the manuscript.

## Authors contributions

AMS, GJF, MH and LH performed all the experiments except for the single cell sequencing. LB, EN and EM performed the single cell sequencing and bioinformatic analysis. GJF, AMS and SM interpreted the data. MR and EM provided intellectual input. AMS and SM wrote the manuscript. GJF, WWS and SM supervised the project, designed the study and provided intellectual, conceptional and scientific advice. All authors critically read the manuscript.

## Supplementary Figures

**Fig. S1.**
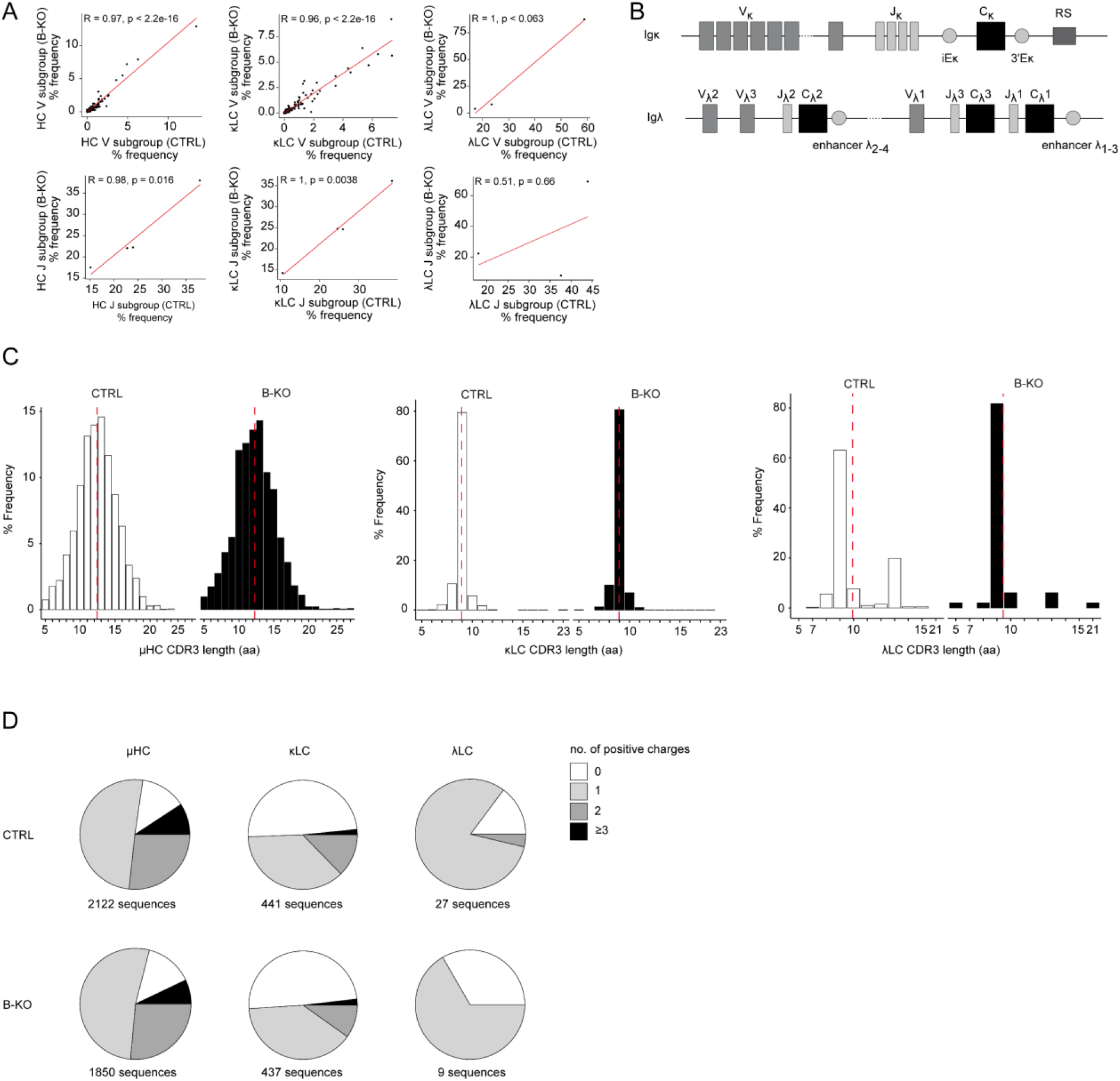
Immature B-KO B cells show a skewed primary BCR repertoire. (**A**, **C**, **D**) Immature B cells (B220^+^IgD^-^IgM^+^) of three individual mice per genotype were pooled and subjected to single cell sequencing analyzing full-length *Ig-gene* V(D)J recombination status and BCR repertoire based on cDNA. More than 10.000 cells per genotype were analyzed. Only productive recombinations are used for the analysis. (**A**) Correlation of *V-* and *J*-gene usage in HC, κLC and λLC between CTRL and B-KO immature B cells. Pearson correlation coefficient (R) and *P* values are shown. Sequence frequencies of the individual *V-* and *J-* subgroup genes of all obtained *V-* and *J*-gene sequences within an individual Ig chain are plotted. (**B**) Schematic representation of the murine *κLC* and *λLC* loci. The CDR3 length (**C**) and amount of positive amino acids (**D**) for HC, κLC and λLC are depicted. Sequence frequencies and distribution within an individual Ig chain are plotted. Mean is represented by a red dotted line (**C**).

**Fig. S2.**
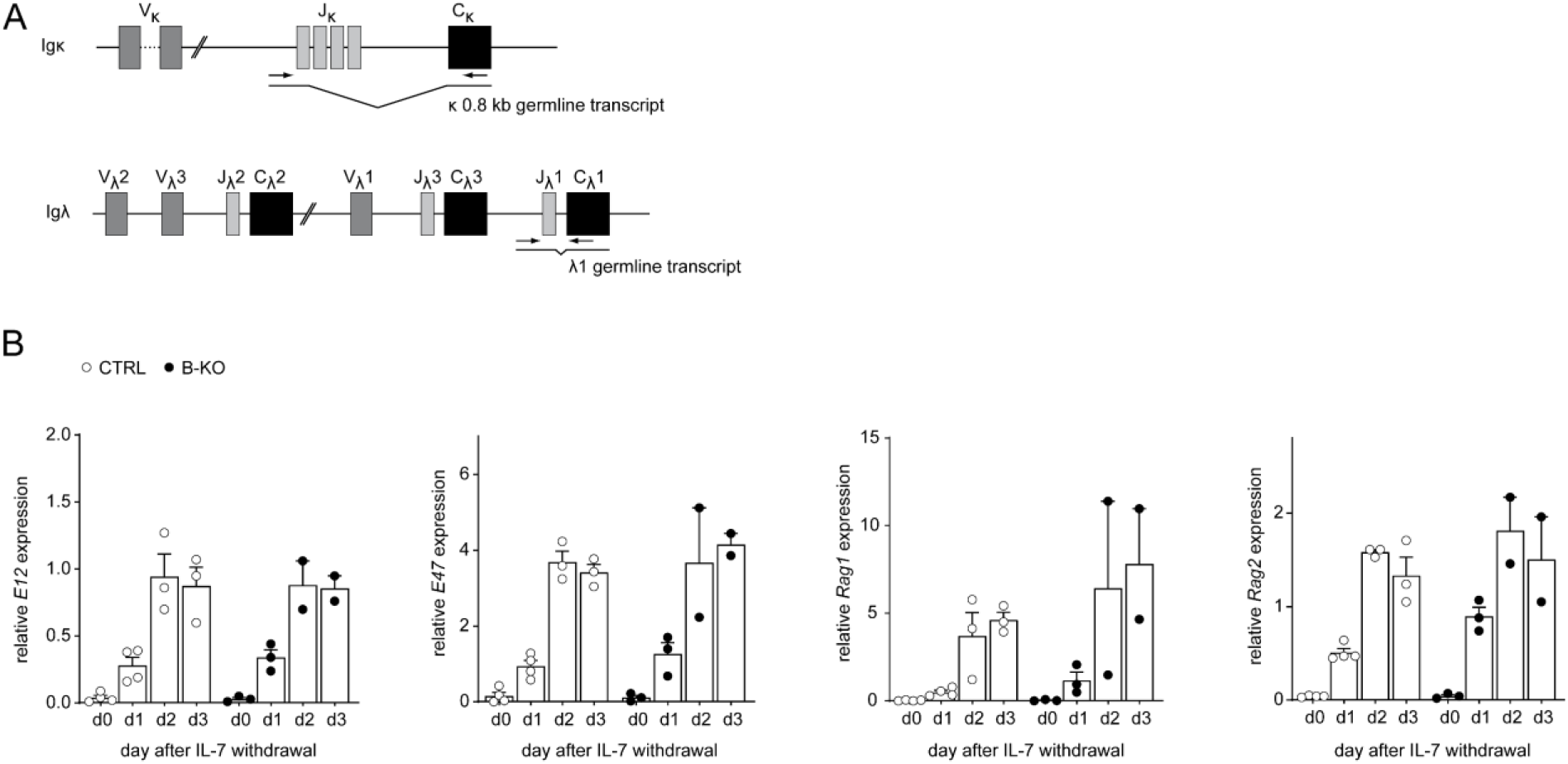
Kidins220 does not regulate *E2A* and *Rag* gene expression. (**A**) Schematic representation of the primer position (black arrows) for the detection of the *Igκ^0^* and *Igλ^1^* germline transcripts. (**B**) RNA isolated from *in vitro* pro-/pre-B cell cultures after IL-7 withdrawal was reverse transcribed to assess the relative expression of mRNA transcripts of *E12*, *E47*, *Rag1* and *Rag2* genes by qRT-PCR. Normalization was done using *Hrpt* (n = 2-4 per genotype; each symbol represents one mouse). In all graphs, the mean ± SEM is plotted. Statistical analysis compared CTRL and B-KO samples for each day using ANOVA test. Only *P* values < 0.05 are indicated.

**Fig. S3.**
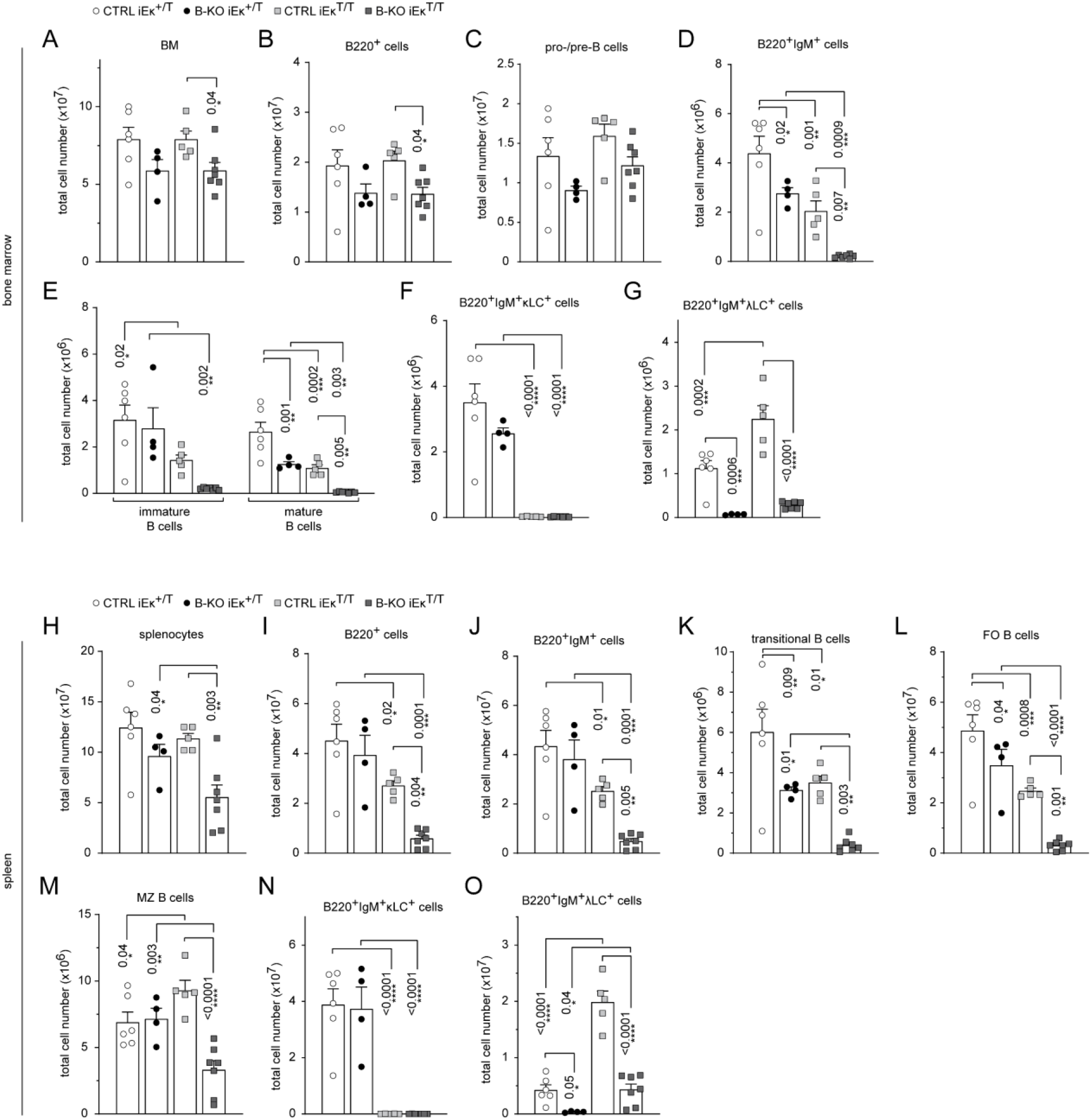
Kidins220 is required for the generation of λLC bearing B cells, even in κ-KO mice. Total cell numbers for each B cell compartment depicted in Figure 3 from BM (**A-G**) and spleen (**H-O**). Pro-/pre-B cells are defined as B220^+^IgM^-^IgD^-^, immature B cells as B220^+^IgM^+^IgD^-^, mature B cells as B220^+^IgM^+^IgD^+^, transitional B cells as B220^+^CD93^+^, follicular (FO) B cells as B220^+^CD93^-^IgM^+^CD21^med^CD23^high^ and marginal zone (MZ) B cells as B220^+^CD93^-^IgM^+^CD21^high^CD23^med^. For the quantification, three independent experiments with n = 4^-^7 mice per genotype were pooled. In all graphs, the mean ± SEM is plotted. Statistical analysis was performed using ANOVA test. CTRL and B-KO in iEκ^+/T^ or iEκ^T/T^ mice were compared. iEκ^+/T^ and iEκ^T/T^ mice were compared for CTRL or B-KO backgrounds. Only *P* values < 0.05 are indicated.

**Fig. S4.**
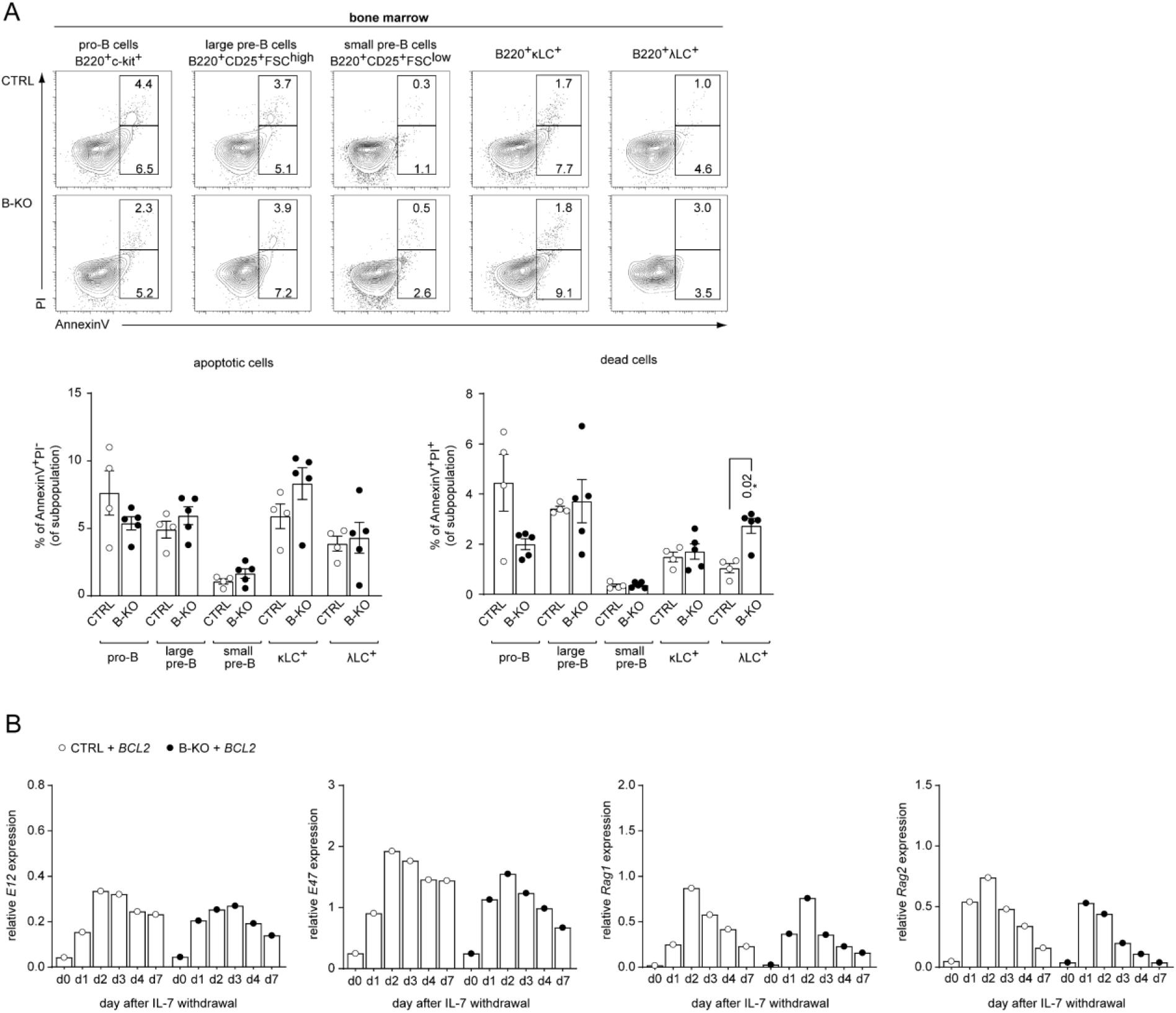
Kidins220 influences B cell survival. (**A**) BM cells were freshly isolated from CTRL and B-KO mice and analyzed for apoptosis and cell viability. The frequency of apoptotic (AnnexinV^+^PI^-^) and dead (AnnexinV^+^PI^+^) cells was analyzed by flow cytometry for B cells at various stages of development in the BM. Representative FACS plots and statistics are depicted (two independent experiments; n = 4-5 mice per genotype). In all graphs, the mean ± SEM is plotted. Statistical analysis was performed using Mann-Whitney test comparing CTRL and B-KO within each population. Only *P* values < 0.05 are indicated. (**B**) RNA was isolated and reverse transcribed to quantify the relative expression of mRNA transcripts of *E12*, *E47*, *Rag1* and *Rag2* genes by qRT-PCR. Data were normalized to *Hrpt* (one mouse per genotype was used).

**Fig. S5.**
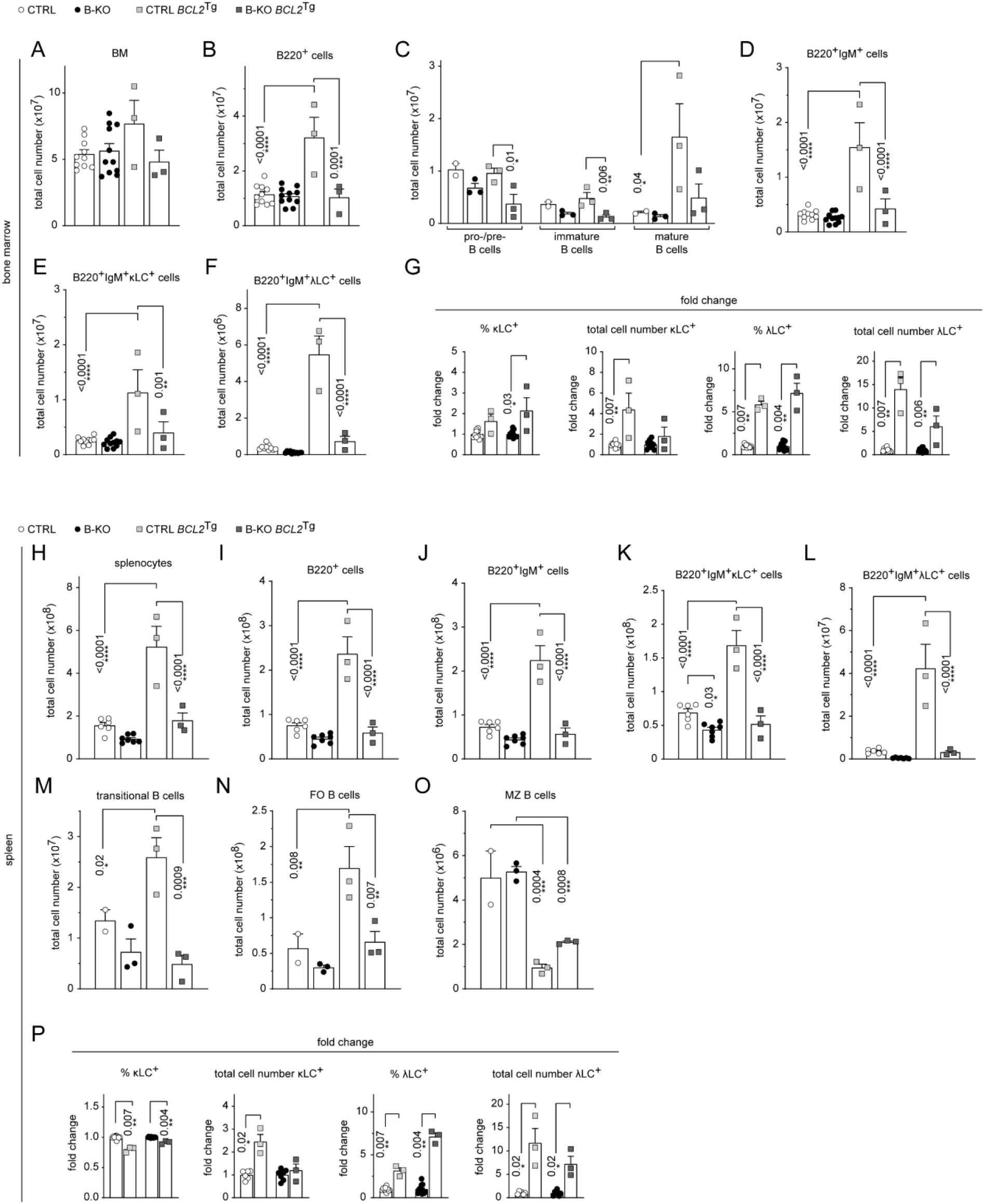
Ectopic BCL2 expression partially rescues λLC expression in B-KO mice. Total cell numbers for each B cell compartment depicted in Figure 5 for the BM (**A-G**) and the spleen (**H-P**) are shown. Pro-/pre-B cells are defined as B220^+^IgM^-^IgD^-^, immature B cells as B220^+^IgM^+^IgD^-^, mature B cells as B220^+^IgM^+^IgD^+^, transitional B cells as B220^+^CD93^+^, follicular (FO) B cells as B220^+^CD93^-^IgM^+^CD21^med^CD23^high^ and marginal zone (MZ) B cells as B220^+^CD93^-^IgM^+^CD21^high^CD23^med^. The quantification of three to five independent experiments with n = 2-12 mice per genotype were pooled. In all graphs, the mean ± SEM is plotted. Statistical analysis was performed using ANOVA test. CTRL and B-KO in the presence or absence of the *BCL2* transgene were compared. Non-transgenic and *BCL2^Tg^* mice were compared for CTRL or B-KO backgrounds. (**O, P**) The fold change of percent and total cell numbers of κLC and λLC B cells is shown in the BM (**O**) and spleen (**P**) in *BCL2^Tg^* relative to non-transgenic mice. The mean ± SEM is plotted. Statistical analysis was performed using Mann-Whitney test. In all graphs, only *P* values < 0.05 are indicated.

**Fig. S6.**
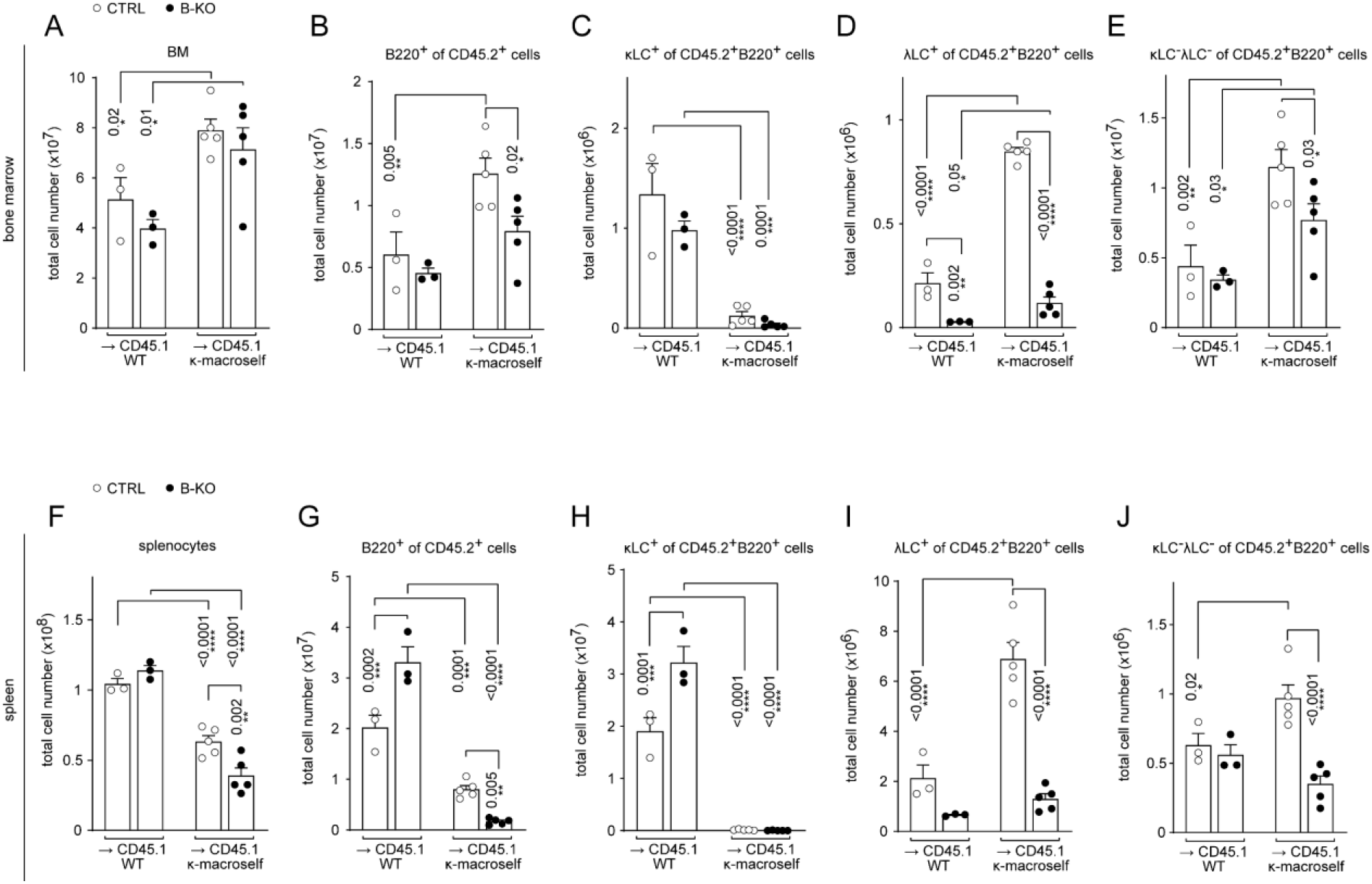
Kidins220 is dispensable for the elimination of autoreactive BCRs but is necessary for the expression of innocuous λLC during tolerance induction. Total cell numbers of each B cell compartment depicted in Figure 6 of BM (**A-E**) and spleen (**F-J**) are shown. The quantification of three independent experiments with n = 6-13 mice per genotype were pooled. In all graphs, the mean ± SEM is plotted. Statistical analysis was performed using ANOVA test. CTRL and B-KO in the presence or absence of the κ-macroself transgene were compared. WT and κ-macroself transgenic mice were compared for CTRL or B-KO backgrounds. Only *P* values < 0.05 are indicated.

**Fig. S7.**
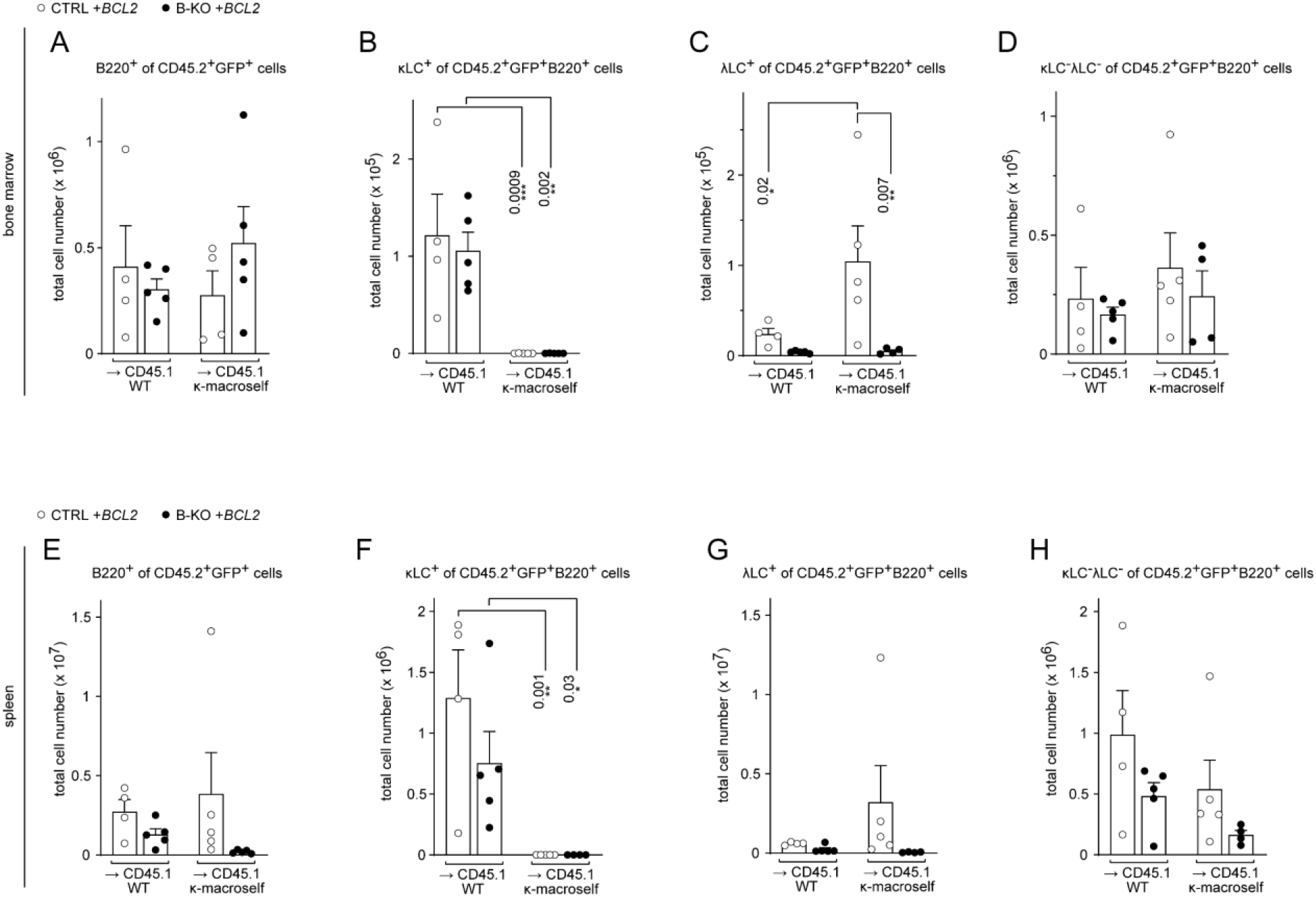
BCL2 overexpression partially rescues λLC expression during tolerance induction. Total cell numbers of each B cell compartment depicted in Figure 7 of BM (**A-D**) and spleen (**E-H**) are shown. BCL2-expressing donor B cells were analyzed by pre-gating on GFP^+^ cells. The quantification of two independent experiments with n = 5-9 mice per genotype were pooled. In all graphs, the mean ± SEM is plotted. Statistical analysis was performed using ANOVA test. CTRL and B-KO in the presence or absence of the κ-macroself transgene were compared. WT and κ-macroself transgenic mice were compared for CTRL or B-KO backgrounds. Only *P* values < 0.05 are indicated.

## References

1. M. R. Clark, M. Mandal, K. Ochiai, H. Singh, Orchestrating B cell lymphopoiesis through interplay of IL-7 receptor and pre-B cell receptor signalling. Nat Rev Immunol. 14, 69–80 (2014), doi:10.1038/nri3570.

2. S. Herzog, M. Reth, H. Jumaa, Regulation of B-cell proliferation and differentiation by pre-B-cell receptor signalling. Nat Rev Immunol. 9, 195–205 (2009), doi:10.1038/nri2491.

3. D. G. Schatz, M. A. Oettinger, D. Baltimore, The V(D)J recombination activating gene, RAG-1. Cell. 59, 1035–1048 (1989), doi:10.1016/0092-8674(89)90760-5.

4. M. A. Oettinger, D. G. Schatz, C. Gorka, D. Baltimore, RAG-1 and RAG-2, adjacent genes that synergistically activate V(D)J recombination. Science (New York, N.Y.). 248, 1517–1523 (1990), doi:10.1126/science.2360047.

5. A. G. Rolink et al., B cell development in the mouse from early progenitors to mature B cells. Immunology Letters. 68, 89–93 (1999), doi:10.1016/S0165-2478(99)00035-8.

6. H. Karasuyama, A. Kudo, F. Melchers, The proteins encoded by the VpreB and lambda 5 pre-B cell-specific genes can associate with each other and with mu heavy chain. Journal of Experimental Medicine. 172, 969–972 (1990), doi:10.1084/jem.172.3.969.

7. T. Tsubata, M. Reth, The products of pre-B cell-specific genes (lambda 5 and VpreB) and the immunoglobulin mu chain form a complex that is transported onto the cell surface. Journal of Experimental Medicine. 172, 973–976 (1990), doi:10.1084/jem.172.3.973.

8. M. G. Reth, P. Ammirati, S. Jackson, F. W. Alt, Regulated progression of a cultured pre-B-cell line to the B-cell stage. Nature. 317, 353–355 (1985), doi:10.1038/317353a0.

9. K. Ochiai et al., A self-reinforcing regulatory network triggered by limiting IL-7 activates pre-BCR signaling and differentiation. Nat Immunol. 13, 300–307 (2012), doi:10.1038/ni.2210.

10. A. Flemming, T. Brummer, M. Reth, H. Jumaa, The adaptor protein SLP-65 acts as a tumor suppressor that limits pre-B cell expansion. Nature immunology. 4, 38–43 (2003), doi:10.1038/ni862.

11. M. Mandal et al., Ras orchestrates exit from the cell cycle and light-chain recombination during early B cell development. Nature immunology. 10, 1110–1117 (2009), doi:10.1038/ni.1785.

12. S. Ma et al., Ikaros and Aiolos inhibit pre-B-cell proliferation by directly suppressing c-Myc expression. Molecular and Cellular Biology. 30, 4149–4158 (2010), doi:10.1128/MCB.00224-10.

13. M. Reth, P. Nielsen, Signaling circuits in early B-cell development. Advances in immunology. 122, 129–175 (2014), doi:10.1016/B978-0-12-800267-4.00004-3.

14. P. Matthias, A. G. Rolink, Transcriptional networks in developing and mature B cells. Nat Rev Immunol. 5, 497–508 (2005), doi:10.1038/nri1633.

15. L. Bai et al., Phospholipase Cgamma2 contributes to light-chain gene activation and receptor editing. Molecular and Cellular Biology. 27, 5957–5967 (2007), doi:10.1128/MCB.02273-06.

16. R. Stadhouders et al., Pre-B cell receptor signaling induces immunoglobulin κ locus accessibility by functional redistribution of enhancer-mediated chromatin interactions. PLoS Biology. 12, e1001791 (2014), doi:10.1371/journal.pbio.1001791.

17. S. Ma, S. Pathak, L. Trinh, R. Lu, Interferon regulatory factors 4 and 8 induce the expression of Ikaros and Aiolos to down-regulate pre-B-cell receptor and promote cell-cycle withdrawal in pre-B-cell development. Blood. 111, 1396–1403 (2008), doi:10.1182/blood-2007-08-110106.

18. R. Lu, K. L. Medina, D. W. Lancki, H. Singh, IRF-4,8 orchestrate the pre-B-to-B transition in lymphocyte development. Genes & development. 17, 1703–1708 (2003), doi:10.1101/gad.1104803.

19. S. Ma, A. Turetsky, L. Trinh, R. Lu, IFN regulatory factor 4 and 8 promote Ig light chain kappa locus activation in pre-B cell development. Journal of immunology (Baltimore, Md.: 1950). 177, 7898–7904 (2006), doi:10.4049/jimmunol.177.11.7898.

20. A. S. Lazorchak, M. S. Schlissel, Y. Zhuang, E2A and IRF-4/Pip promote chromatin modification and transcription of the immunoglobulin kappa locus in pre-B cells. Molecular and Cellular Biology. 26, 810–821 (2006), doi:10.1128/MCB.26.3.810-821.2006.

21. K. Beck, M. M. Peak, T. Ota, D. Nemazee, C. Murre, Distinct roles for E12 and E47 in B cell specification and the sequential rearrangement of immunoglobulin light chain loci. The Journal of experimental medicine. 206, 2271–2284 (2009), doi:10.1084/jem.20090756.

22. S. Pathak, S. Ma, L. Trinh, R. Lu, A role for interferon regulatory factor 4 in receptor editing. Molecular and Cellular Biology. 28, 2815–2824 (2008), doi:10.1128/MCB.01946-07.

23. M. A. Inlay, H. Tian, T. Lin, Y. Xu, Important roles for E protein binding sites within the immunoglobulin kappa chain intronic enhancer in activating Vkappa Jkappa rearrangement. The Journal of experimental medicine. 200, 1205–1211 (2004), doi:10.1084/jem.20041135.

24. K. Johnson et al., Regulation of immunoglobulin light-chain recombination by the transcription factor IRF-4 and the attenuation of interleukin-7 signaling. Immunity. 28, 335–345 (2008), doi:10.1016/j.immuni.2007.12.019.

25. M. W. Quong et al., Receptor editing and marginal zone B cell development are regulated by the helix-loop-helix protein, E2A. The Journal of experimental medicine. 199, 1101–1112 (2004), doi:10.1084/jem.20031180.

26. R. Kersseboom et al., Bruton’s tyrosine kinase and SLP-65 regulate pre-B cell differentiation and the induction of Ig light chain gene rearrangement. Journal of immunology (Baltimore, Md.: 1950). 176, 4543–4552 (2006), doi:10.4049/jimmunol.176.8.4543.

27. G. M. Dingjan et al., Bruton’s tyrosine kinase regulates the activation of gene rearrangements at the lambda light chain locus in precursor B cells in the mouse. The Journal of experimental medicine. 193, 1169–1178 (2001), doi:10.1084/jem.193.10.1169.

28. K. Hayashi, T. Nojima, R. Goitsuka, D. Kitamura, Impaired receptor editing in the primary B cell repertoire of BASH-deficient mice. Journal of immunology (Baltimore, Md.: 1950). 173, 5980–5988 (2004), doi:10.4049/jimmunol.173.10.5980.

29. S. Xu, K.-G. Lee, J. Huo, T. Kurosaki, K.-P. Lam, Combined deficiencies in Bruton tyrosine kinase and phospholipase Cgamma2 arrest B-cell development at a pre-BCR+ stage. Blood. 109, 3377–3384 (2007), doi:10.1182/blood-2006-07-036418.

30. A. Hashimoto et al., Cutting edge: essential role of phospholipase C-gamma 2 in B cell development and function. Journal of immunology (Baltimore, Md.: 1950). 165, 1738–1742 (2000), doi:10.4049/jimmunol.165.4.1738.

31. Y. Sun, Z. Wei, N. Li, Y. Zhao, A comparative overview of immunoglobulin genes and the generation of their diversity in tetrapods. Developmental and comparative immunology. 39, 103–109 (2013), doi:10.1016/j.dci.2012.02.008.

32. K. L. McGuire, E. S. Vitetta, kappa/lambda Shifts do not occur during maturation of murine B cells. Journal of immunology (Baltimore, Md.: 1950). 127, 1670–1673 (1981).

33. D. Nemazee, Receptor editing in lymphocyte development and central tolerance. Nat Rev Immunol. 6, 728–740 (2006), doi:10.1038/nri1939.

34. E. T. Luning Prak, M. Monestier, R. A. Eisenberg, B cell receptor editing in tolerance and autoimmunity. Annals of the New York Academy of Sciences. 1217, 96–121 (2011), doi:10.1111/j.1749-6632.2010.05877.x.

35. D. Ait-Azzouzene et al., An immunoglobulin C kappa-reactive single chain antibody fusion protein induces tolerance through receptor editing in a normal polyclonal immune system. The Journal of experimental medicine. 201, 817–828 (2005), doi:10.1084/jem.20041854.

36. E. Derudder et al., Development of immunoglobulin lambda-chain-positive B cells, but not editing of immunoglobulin kappa-chain, depends on NF-kappaB signals. Nature immunology. 10, 647–654 (2009), doi:10.1038/ni.1732.

37. G. J. Fiala et al., Kidins220/ARMS binds to the B cell antigen receptor and regulates B cell development and activation. The Journal of experimental medicine. 212, 1693–1708 (2015), doi:10.1084/jem.20141271.

38. T. Iglesias et al., Identification and cloning of Kidins220, a novel neuronal substrate of protein kinase D. Journal of Biological Chemistry. 275, 40048–40056 (2000), doi:10.1074/jbc.M005261200.

39. H. Kong, J. Boulter, J. L. Weber, C. Lai, M. V. Chao, An Evolutionarily Conserved Transmembrane Protein That Is a Novel Downstream Target of Neurotrophin and Ephrin Receptors. J. Neurosci. 21, 176–185 (2001), doi:10.1523/JNEUROSCI.21-01-00176.2001.

40. S. Deswal et al., Kidins220/ARMS associates with B-Raf and the TCR, promoting sustained Erk signaling in T cells. Journal of immunology (Baltimore, Md.: 1950). 190, 1927–1935 (2013), doi:10.4049/jimmunol.1200653.

41. V. E. Neubrand, F. Cesca, F. Benfenati, G. Schiavo, Kidins220/ARMS as a functional mediator of multiple receptor signalling pathways. Journal of Cell Science. 125, 1845–1854 (2012), doi:10.1242/jcs.102764.

42. F. Cesca et al., Kidins220/ARMS mediates the integration of the neurotrophin and VEGF pathways in the vascular and nervous systems. Cell Death Differ. 19, 194–208 (2012), doi:10.1038/cdd.2011.141.

43. A. L. Kenter, C. T. Watson, J.-H. Spille, Igh Locus Polymorphism May Dictate Topological Chromatin Conformation and V Gene Usage in the Ig Repertoire. Front. Immunol. 12, 682589 (2021), doi:10.3389/fimmu.2021.682589.

44. M. Aoki-Ota, A. Torkamani, T. Ota, N. Schork, D. Nemazee, Skewed primary Igκ repertoire and V-J joining in C57BL/6 mice: implications for recombination accessibility and receptor editing. Journal of immunology (Baltimore, Md.: 1950). 188, 2305–2315 (2012), doi:10.4049/jimmunol.1103484.

45. E. L. Prak, M. Trounstine, D. Huszar, M. Weigert, Light chain editing in kappa-deficient animals: a potential mechanism of B cell tolerance. The Journal of experimental medicine. 180, 1805–1815 (1994), doi:10.1084/jem.180.5.1805.

46. M. W. Retter, D. Nemazee, Receptor editing occurs frequently during normal B cell development. The Journal of experimental medicine. 188, 1231–1238 (1998), doi:10.1084/jem.188.7.1231.

47. T. Gerdes, M. Wabl, Physical map of the mouse lambda light chain and related loci. Immunogenetics. 54, 62–65 (2002), doi:10.1007/s00251-002-0435-y.

48. P. Sanchez, D. Rueff-Juy, P. Boudinot, S. Hachemi-Rachedi, P. A. Cazenave, The lambda B cell repertoire of kappa-deficient mice. International Reviews of Immunology. 13, 357–368 (1996), doi:10.3109/08830189609061758.

49. P. Boudinot, A. M. Drapier, P. A. Cazenave, P. Sanchez, Conserved distribution of lambda subtypes from rearranged gene segments to immunoglobulin synthesis in the mouse B cell repertoire. Eur. J. Immunol. 24, 2013–2017 (1994), doi:10.1002/eji.1830240912.

50. P. Sanchez, B. Nadel, P. A. Cazenave, V lambda-J lambda rearrangements are restricted within a V-J-C recombination unit in the mouse. Eur. J. Immunol. 21, 907–911 (1991), doi:10.1002/eji.1830210408.

51. E. B. Reilly, B. Blomberg, T. Imanishi-Kari, S. Tonegawa, H. N. Eisen, Restricted association of V and J-C gene segments for mouse lambda chains. Proceedings of the National Academy of Sciences of the United States of America. 81, 2484–2488 (1984), doi:10.1073/pnas.81.8.2484.

52. E. Miho, R. Roškar, V. Greiff, S. T. Reddy, Large-scale network analysis reveals the sequence space architecture of antibody repertoires. Nature communications. 10, 1321 (2019), doi:10.1038/s41467-019-09278-8.

53. A. Q. Maranhão et al., A mouse variable gene fragment binds to DNA independently of the BCR context: a possible role for immature B-cell repertoire establishment. PloS one. 8, e72625 (2013), doi:10.1371/journal.pone.0072625.

54. M. A. Zelazowska et al., Gammaherpesvirus-infected germinal center cells express a distinct immunoglobulin repertoire. Life science alliance. 3 (2020), doi:10.26508/lsa.201900526.

55. E. Smakaj et al., Benchmarking immunoinformatic tools for the analysis of antibody repertoire sequences. Bioinformatics. 36, 1731–1739 (2020), doi:10.1093/bioinformatics/btz845.

56. A. Rolink, M. Streb, F. Melchers, The kappa/lambda ratio in surface immunoglobulin molecules on B lymphocytes differentiating from DHJH-rearranged murine pre-B cell clones in vitro. Eur. J. Immunol. 21, 2895–2898 (1991), doi:10.1002/eji.1830211137.

57. S. Middendorp, G. M. Dingjan, R. W. Hendriks, Impaired precursor B cell differentiation in Bruton’s tyrosine kinase-deficient mice. Journal of immunology (Baltimore, Md.: 1950). 168, 2695–2703 (2002), doi:10.4049/jimmunol.168.6.2695.

58. M. S. Schlissel, D. Baltimore, Activation of immunoglobulin kappa gene rearrangement correlates with induction of germline kappa gene transcription. Cell. 58, 1001–1007 (1989), doi:10.1016/0092-8674(89)90951-3.

59. H. Engel, A. Rolink, S. Weiss, B cells are programmed to activate κ and λ for rearrangement at consecutive developmental stages. Eur. J. Immunol. 29, 2167–2176 (1999), doi:10.1002/(SICI)1521-4141(199907)29:07<2167::AID-IMMU2167>3.0.CO;2-H.

60. H. Engel, H. Rühl, C. J. Benham, J. Bode, S. Weiss, Germ-line transcripts of the immunoglobulin λ J–C clusters in the mouse: characterization of the initiation sites and regulatory elements. Molecular Immunology. 38, 289–302 (2001), doi:10.1016/S0161-5890(01)00056-6.

61. L.-Y. Hsu et al., A Conserved Transcriptional Enhancer Regulates RAG Gene Expression in Developing B Cells. Immunity. 19, 105–117 (2003), doi:10.1016/S1074-7613(03)00181-X.

62. S. Takeda et al., Deletion of the immunoglobulin kappa chain intron enhancer abolishes kappa chain gene rearrangement in cis but not lambda chain gene rearrangement in trans. The EMBO journal. 12, 2329–2336 (1993).

63. S. Pillai, A. Cariappa, The follicular versus marginal zone B lymphocyte cell fate decision. Nat Rev Immunol. 9, 767–777 (2009), doi:10.1038/nri2656.

64. S. Ogilvy et al., Constitutive Bcl-2 expression throughout the hematopoietic compartment affects multiple lineages and enhances progenitor cell survival. Proceedings of the National Academy of Sciences of the United States of America. 96, 14943–14948 (1999), doi:10.1073/pnas.96.26.14943.

65. M. Yabas et al., ATP11C is critical for the internalization of phosphatidylserine and differentiation of B lymphocytes. Nature immunology. 12, 441–449 (2011), doi:10.1038/ni.2011.

66. C. J. Vandenberg et al., Loss of Bak enhances lymphocytosis but does not ameliorate thrombocytopaenia in BCL-2 transgenic mice. Cell Death Differ. 21, 676–684 (2014), doi:10.1038/cdd.2013.201.

67. A. Banerjee et al., NF-kappaB1 and c-Rel cooperate to promote the survival of TLR4-activated B cells by neutralizing Bim via distinct mechanisms. Blood. 112, 5063–5073 (2008), doi:10.1182/blood-2007-10-120832.

68. E. Derudder et al., Canonical NF-κB signaling is uniquely required for the long-term persistence of functional mature B cells. Proceedings of the National Academy of Sciences of the United States of America. 113, 5065–5070 (2016), doi:10.1073/pnas.1604529113.

69. J. Lang et al., Enforced Bcl-2 expression inhibits antigen-mediated clonal elimination of peripheral B cells in an antigen dose-dependent manner and promotes receptor editing in autoreactive, immature B cells. The Journal of experimental medicine. 186, 1513–1522 (1997), doi:10.1084/jem.186.9.1513.

70. J. L. Vela, D. Aït-Azzouzene, B. H. Duong, T. Ota, D. Nemazee, Rearrangement of mouse immunoglobulin kappa deleting element recombining sequence promotes immune tolerance and lambda B cell production. Immunity. 28, 161–170 (2008), doi:10.1016/j.immuni.2007.12.011.

71. R. Kersseboom et al., Bruton’s tyrosine kinase cooperates with the B cell linker protein SLP-65 as a tumor suppressor in Pre-B cells. Journal of Experimental Medicine. 198, 91–98 (2003), doi:10.1084/jem.20030615.

72. T. Tsubata, Involvement of Reactive Oxygen Species (ROS) in BCR Signaling as a Second Messenger. Advances in experimental medicine and biology. 1254, 37–46 (2020), doi:10.1007/978-981-15-3532-1_3.

73. M. Stein et al., A defined metabolic state in pre B cells governs B-cell development and is counterbalanced by Swiprosin-2/EFhd1. Cell death and differentiation. 24, 1239–1252 (2017), doi:10.1038/cdd.2017.52.

74. S. F. Haque, S. L. Bevington, J. Boyes, The Eλ3-1 enhancer is essential for V(D)J recombination of the murine immunoglobulin lambda light chain locus. Biochemical and Biophysical Research Communications. 441, 482–487 (2013), doi:10.1016/j.bbrc.2013.10.087.

75. A. M. Collins, C. T. Watson, Immunoglobulin Light Chain Gene Rearrangements, Receptor Editing and the Development of a Self-Tolerant Antibody Repertoire. Front. Immunol. 9, 2249 (2018), doi:10.3389/fimmu.2018.02249.

76. J. C. Arévalo, H. Yano, K. K. Teng, M. V. Chao, A unique pathway for sustained neurotrophin signaling through an ankyrin-rich membrane-spanning protein. The EMBO journal. 23, 2358–2368 (2004), doi:10.1038/sj.emboj.7600253.

77. C. López-Menéndez et al., Kidins220/ARMS downregulation by excitotoxic activation of NMDARs reveals its involvement in neuronal survival and death pathways. Journal of Cell Science. 122, 3554–3565 (2009), doi:10.1242/jcs.056473.

78. K. Zhang et al., SINO Syndrome Causative KIDINS220/ARMS Gene Regulates Adipocyte Differentiation. Front. Cell Dev. Biol. 9, 619475 (2021), doi:10.3389/fcell.2021.619475.

79. F. Jaudon et al., Kidins220/ARMS controls astrocyte calcium signaling and neuron-astrocyte communication. Cell Death Differ. 27, 1505–1519 (2020), doi:10.1038/s41418-019-0431-5.

80. F. Jaudon, M. Albini, S. Ferroni, F. Benfenati, F. Cesca, A developmental stage-and Kidins220-dependent switch in astrocyte responsiveness to brain-derived neurotrophic factor. Journal of Cell Science. 134 (2021), doi:10.1242/jcs.258419.

81. J.-W. Lee et al., CD25 (IL2RA) Orchestrates Negative Feedback Control and Stabilizes Oncogenic Signaling Strength in Acute Lymphoblastic Leukemia. Blood. 126, 1434 (2015), doi:10.1182/blood.V126.23.1434.1434.

82. A. Rolink, U. Grawunder, T. H. Winkler, H. Karasuyama, F. Melchers, IL-2 receptor alpha chain (CD25, TAC) expression defines a crucial stage in pre-B cell development. Int Immunol. 6, 1257–1264 (1994), doi:10.1093/intimm/6.8.1257.

83. Y.-Y. Feng et al., Essential Role of NADPH Oxidase-Dependent Production of Reactive Oxygen Species in Maintenance of Sustained B Cell Receptor Signaling and B Cell Proliferation. Journal of immunology (Baltimore, Md.: 1950). 202, 2546–2557 (2019), doi:10.4049/jimmunol.1800443.

84. M. L. Wheeler, A. L. Defranco, Prolonged production of reactive oxygen species in response to B cell receptor stimulation promotes B cell activation and proliferation. Journal of immunology (Baltimore, Md.: 1950). 189, 4405–4416 (2012), doi:10.4049/jimmunol.1201433.

85. E. Meffre, M. C. Nussenzweig, Deletion of immunoglobulin beta in developing B cells leads to cell death. Proceedings of the National Academy of Sciences of the United States of America. 99, 11334–11339 (2002), doi:10.1073/pnas.172369999.

86. M. Kraus, M. B. Alimzhanov, N. Rajewsky, K. Rajewsky, Survival of resting mature B lymphocytes depends on BCR signaling via the Igalpha/beta heterodimer. Cell. 117, 787–800 (2004), doi:10.1016/j.cell.2004.05.014.

87. T. Yasuda et al., Erk kinases link pre-B cell receptor signaling to transcriptional events required for early B cell expansion. Immunity. 28, 499–508 (2008), doi:10.1016/j.immuni.2008.02.015.

88. U. Siebenlist, K. Brown, E. Claudio, Control of lymphocyte development by nuclear factor-kappaB. Nature reviews. Immunology. 5, 435–445 (2005), doi:10.1038/nri1629.

89. J. J. Bednarski et al., RAG-induced DNA double-strand breaks signal through Pim2 to promote pre-B cell survival and limit proliferation. Journal of Experimental Medicine. 209, 11–17 (2012), doi:10.1084/jem.20112078.

90. L. A. O’Reilly et al., MEK/ERK-mediated phosphorylation of Bim is required to ensure survival of T and B lymphocytes during mitogenic stimulation. Journal of immunology (Baltimore, Md.: 1950). 183, 261–269 (2009), doi:10.4049/jimmunol.0803853.

91. M. R. Gold, B cell development: important work for ERK. Immunity. 28, 488–490 (2008), doi:10.1016/j.immuni.2008.03.008.

92. H. Wardemann, J. Hammersen, M. C. Nussenzweig, Human autoantibody silencing by immunoglobulin light chains. Journal of Experimental Medicine. 200, 191–199 (2004), doi:10.1084/jem.20040818.

93. C. L. Townsend et al., Significant Differences in Physicochemical Properties of Human Immunoglobulin Kappa and Lambda CDR3 Regions. Front. Immunol. 7, 388 (2016), doi:10.3389/fimmu.2016.00388.

94. R. Novak, E. Jacob, J. Haimovich, O. Avni, D. Melamed, The MAPK/ERK and PI3K pathways additively coordinate the transcription of recombination-activating genes in B lineage cells. Journal of immunology (Baltimore, Md.: 1950). 185, 3239–3247 (2010), doi:10.4049/jimmunol.1001430.

95. L. Verkoczy et al., A role for nuclear factor kappa B/rel transcription factors in the regulation of the recombinase activator genes. Immunity. 22, 519–531 (2005), doi:10.1016/j.immuni.2005.03.006.

96. K. Johnson et al., IL-7 functionally segregates the pro-B cell stage by regulating transcription of recombination mediators across cell cycle. Journal of immunology (Baltimore, Md.: 1950). 188, 6084–6092 (2012), doi:10.4049/jimmunol.1200368.

